# Intra-Ramanome Correlation Analysis Unveils Metabolite Conversion Network from an Isogenic Cellular Population

**DOI:** 10.1101/2020.12.22.424060

**Authors:** Yuehui He, Shi Huang, Peng Zhang, Yuetong Ji, Jian Xu

**Author notes:** Corresponding to: Jian Xu.

## Abstract

Revealing dynamic features of cellular systems, such as links among metabolic phenotypes, typically requires a time- or condition-series set of samples. Here Intra-Ramanome Correlation analysis (IRCA) was proposed to achieve this goal from just one snapshot of an isogenic population, by pairwise correlating among cells all the thousands of Raman bands from Single-cell Raman Spectra (SCRS), i.e., based on the intrinsic inter-cellular metabolic heterogeneity. IRCA of *Chlamydomonas reinhardtii* under nitrogen depletion revealed a metabolite conversion network at each time point and its temporal dynamics that feature protein-to-starch conversion followed by starch-to-TAG conversion (plus conversion of membrane lipids to TAG). Such correlation patterns in IRCA were abrased by knocking out the starch-biosynthesis pathway yet fully restored by genetic complementation. Extension to 64 ramanomes from microalgae, fungi and bacteria under various conditions suggests IRCA-derived metabolite conversion network as an intrinsic, reliable, species-resolved and state-sensitive metabolic signature of isogenic cellular population. The high throughput, low cost, excellent scalability and broad extendibility of IRCA suggest its broad application in cellular systems.

## Introduction

The link, or interaction, among metabolic phenotypes is a fundamental feature that underlies proper function of cellular systems ^1, 2, 3^. As extraction of such information requires variation of metabolic phenotypes, typically a time-series or condition-series set of samples are characterized respectively and then correlated (resolution detecting inter-phenotype links is usually dependent on sample size) ^4, 5, 6^. However, within a single sample of an isogenic population of cells, at any given time or condition, intercellular phenotypic variation is universal and inherent. Can such phenotypic variation among individual cells, instead of that among samples, be exploited to predict the link or interaction among metabolic phenotypes? This hypothesis is built on: (*i*) each cell carries an array of multiple metabolic phenotypes; (*ii*) every cell can be quantitatively phenotyped as one independent biological replicate; (*iii*) inter-cellular phenotypic variation can be measured rapidly, for many cells; (*iv*) inter-cellular correlation of the metabolic phenotypes can unveil important features of the system.

To test this hypothesis, we employed Single-cell Raman Spectra (SCRS), which captures the *in vivo* chemical profiles of a cell in a rapid, label-free and non-destructive manner ^7, 8, 9^, with each of its nearly 1600 Raman peaks potentially representing a specific metabolic phenotype (e.g., utilization of a substrate or production of a compound). SCRS, one from a cell, can be randomly sampled from a given instance of an isogenic cellular population, and collectively form a ramanome ^10, 11^ that represents a single-cell-resolution metabolic snapshot of the population. By treating every SCRS as one biological replicate and each of its bands or combination of bands as a metabolic phenotype, here we propose a formal framework called Intra-Ramanome Correlation Analysis (IRCA; **Fig. 1**). From just a single ramanome, by pairwise correlating all Raman bands of SCRS among the individual cells, IRCA unveils a network of potential metabolite conversions. Thus IRCA is a non-invasive, rapid, low-cost, high-throughput, landscape-like, and universally applicable method for unraveling metabolic features of cellular systems.

**Figure 1.**
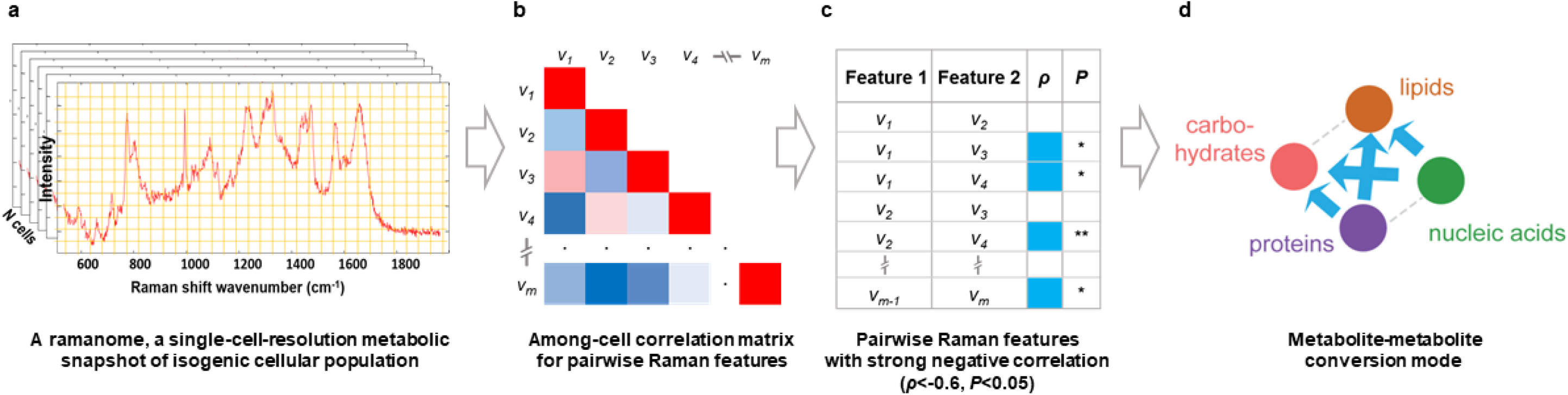
The principle and workflow of Intra-Ramanome Correlation Analysis (IRCA).

## Results

### IRCA predicts inter-conversion among starch, protein and TAG from one instance of population

To test the IRCA concept (**Fig. 1**), we employed as a first model the unicellular alga of *Chlamydomonas reinhardtii* (*Cr*) CC124, which was grown under nitrogen depletion (N-; **Methods**). Triplicate isogenic cultures were sampled from 16 time points from 0h to 8d (48 ramanomes in total; **Fig. S1**). For cellular protein, starch and TAG contents, each sample was analyzed at the population level via various techniques that all require extraction from cell lysates (**Fig. 2a, b, c**; **Supplemental Information**), and at the single-cell level via quantitative models (Partial Least Square Regression, PLSR) based on full SCRS, which is non-invasive and label-free ^12^ (20 randomly selected cells per sample thus 60 per time point; **Fig. 2d, e, f**). Correlation coefficients (R^2^) of protein, starch and TAG contents between the two approaches are 0.9924, 0.9892 and 0.9686 respectively, confirming accuracy of simultaneous quantification for a *Cr* cell via its full SCRS ^12^. Notably, for each phenotype, at any instance of the population, the degree of intercellular heterogeneity is high, and can vary greatly along the eight days (**Fig. S1g, h, i**). Specifically, for protein, both minimal and maximal single-cell contents in the population were decreasing (**Fig. S1g**), while for starch, the trends are exactly opposite (**Fig. S1h**). As for TAG, the maximal content was increasing while the minimal remained at the baseline (i.e., a subpopulation of non-TAG-producing cells was always present), with degree of within-population heterogeneity continuing to grow (i.e., ‘delta’; **Fig. S1i**).

**Figure 2.**
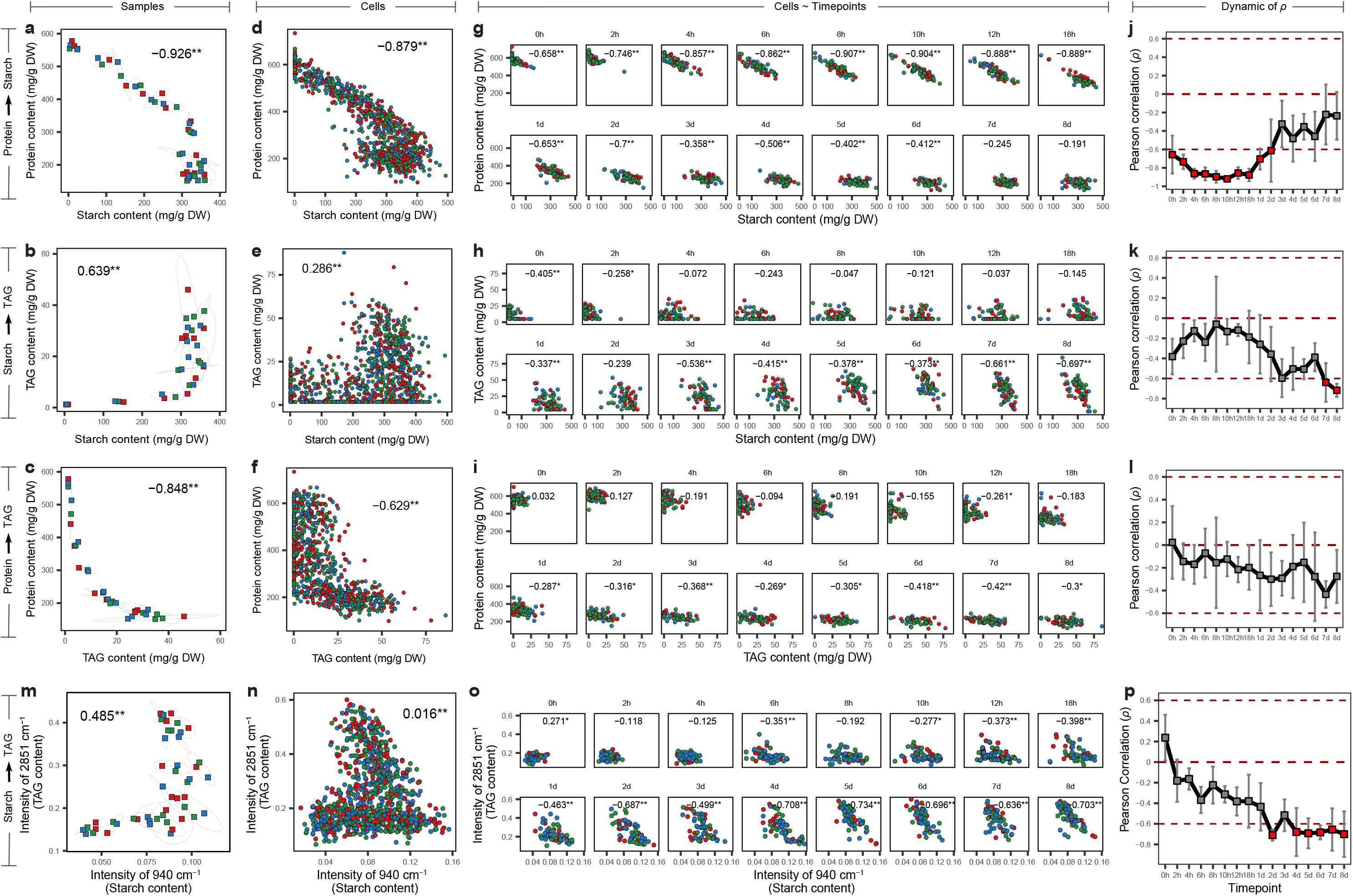
Correlation of starch and TAG content of CC124 at population level and single-cell level. (**a-f**) Correlation of starch and total protein content at population level (**a**) and single-cell level (**d**). Correlation of starch and TAG content at population level (**b**) and single-cell level (**e**). Correlation of TAG and total protein content at population level (**c**) and single-cell level (**f**). Each dot in **a**, **b** and **c** represents one sample. Each dot in **d**, **e** and **f** represents one individual cell. (**g-l**) Correlation of starch and TAG content modeled by multiple pairs of singular Raman bands among individual cells at each time point. Phenotypic correlation between starch (*x*) and protein (*y*) contents (**g**, **j**), between starch (*x*) and TAG (*y*) contents (**h**, **k**), between TAG (*x*) and protein (*y*) contents (**i**, **l**) were shown. (**m**-**p**) Phenotypic correlation between starch (*x*) and TAG (*y*) contents modeled by pair of Raman band of 940 cm-1 and 2851 cm-1. Correlation of starch and TAG content at population level (**m**) and single-cell level (**n**). Phenotypic correlation between starch (*x*) and TAG (*y*) contents (**o**, **p**) at each timepoints were shown. In the temporal dynamics curves, *ρ* is Pearson correlation coefficients among the phenotypes among single cells (**: *P* < 0.01; *: *P* < 0.05), with value indicating mean of triplicates and error bar standard deviation. Absence of correlation (*ρ* = 0) or presence of strong correlation (*ρ* higher than 0.6 or lower than −0.6) was highlighted by red horizontal lines.

At the population level, significant correlation (defined by Pearson correlation coefficient or *ρ*) was observed in each of three phenotype pairs among the 48 populations: protein-starch (*ρ* = −0.926; **Fig. 2a**), starch-TAG (*ρ* = 0.639; **Fig. 3b**) and protein-TAG (*ρ* = −0.848; **Fig. 2c**). This indicates protein-starch conversion, protein-TAG conversion and accordant change of starch and TAG contents ^13, 14^. Correlation at the single-cell level for the 960 cells collectively reached a consistent conclusion, albeit with lower correlation coefficients (*ρ* = −0.879, 0.286 and −0.229 respectively; **Fig. 2d, e, f**). Thus inter-phenotype correlation among pooled cells from all populations can recapitulate that among populations.

**Figure 3.**
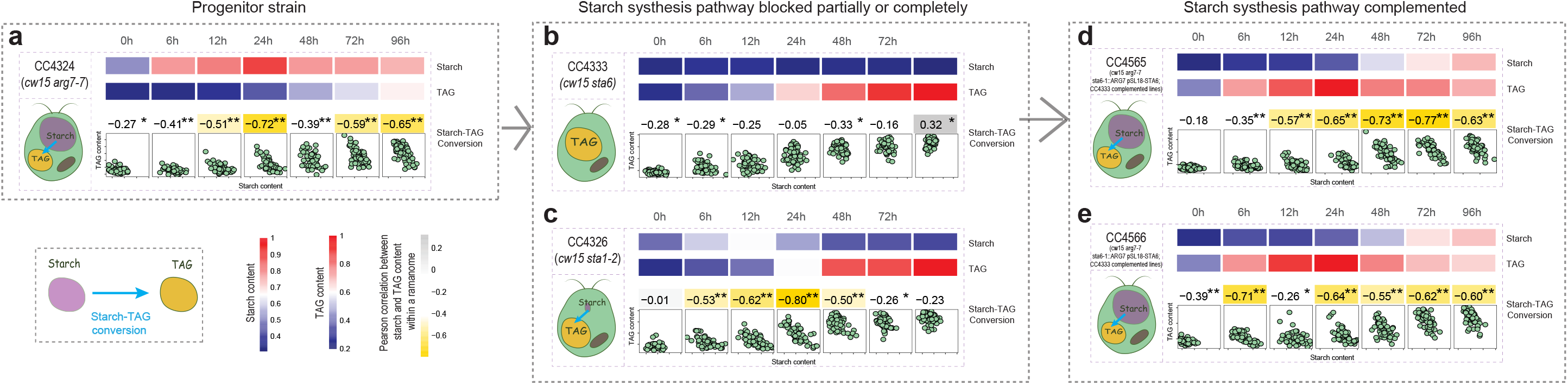
Validation of the IRCA method via a genetic approach, using a series of targeted mutants of *C. reinhardtii*. Correlation between starch (*x*) and TAG (*y*) contents (modeled by 940 cm-1 and 2851 cm-1 respectively) among cells within a ramanome for mutants strains (**Table S1**): CC4324 (**a**), CC4333 (**b**), CC4326 (**c**), CC4565 (**d**), CC4566 (**e**). The starch and TAG contents and Pearson correlation coefficients were shown via heatmap (**: *P* < 0.01; *: *P* < 0.05). Each dot in the scatterplots represents one cell.

We then probed whether such inter-phenotype correlation can be detected via cells from just one instance of population, i.e., at each of the 16 time points (**Fig. 2g, h, i**). Intriguingly, for protein-starch, negative correlation (NC) is phase-specific and weakening: strong at each of the time points only before 2d (*ρ* < −0.6, *P* < 0.01; **Fig. 2g**) and absent at 7d or 8d (**Fig. 2j**). Thus it is possible that protein-starch conversion took place only at the early phase of N-. In fact, this prediction is supported by routing of hydrolytic release of carbon skeletons from protein to starch-granule synthesis during the first two days in *Cr* under N-^13, 14^.

For starch-TAG, the situation was opposite, as the trend of NC among cells within a population intensified (**Fig. 2h, k**). Thus, in contrast to the positive correlation (PC) between starch and TAG at the population level (*ρ* = 0.639, *P* < 0.01; **Fig. 2b**), the within-population NC between starch and TAG was prominently present at the late phase (i.e., starting from 3d; **Fig. 2k**), which predicts conversion between starch and TAG inside cell. This IRCA-based prediction, which the population-level analysis missed, is actually supported by (*i*) competition between starch and lipid synthesis for shared biosynthetic precursors intensifies temporally ^15, 16, 17, 18^, and (*ii*) early build-up of starch may serve as carbon source for lipid synthesis at a later subsequent phase ^19, 20, 21, 22, 23^.

For protein-TAG, unlike their very strong among-population NC (*ρ* = −0.848, *P* < 0.01; **Fig. 2c**), no strong correlation was found for each population, although the trend of NC gradually intensified (**Fig. 2i, l**). In particular, at 2d and beyond, protein contents of individual cells were all already depleted to a very low level, despite their relatively wide range of TAG content, an indication of the decoupling of protein-synthetic and TAG-synthetic pathways. This distinction in temporal within-population NC pattern between protein-starch (**Fig. 2j**) and protein-TAG (**Fig. 2l**) suggests that, (*i*) prior to 2d, majority of released carbon skeletons from proteins were routed to starch rather than lipids; (*ii*) when starch biosynthesis was saturated at 2d, substantial accumulation of TAG occurred which is converted partially from protein ^14^. Therefore IRCA, by inter-phenotype correlation among cells, can reveal inter-metabolite conversions from just one instance of an isogenic population.

Notably, besides the full *Cr* SCRS ^12^, individual Raman bands can also quantitatively model singe-cell starch, protein and TAG contents. For example, 940 cm^−1^ (C-O stretching, C-O-C, C-O-H deformation and α-helix C-C backbone) and 2851 cm^−1^ (C-H_2_, C-H_3_ asymmetric and symmetric stretches) can model starch and TAG respectively (correlation coefficient R^2^ between conventional, bulk-biomass analyses and SCRS-based analyses being 0.884 and 0.954 respectively; **Fig. S2**). Moreover, correlations via just these two peaks among populations (**Fig. 2m**) or among cells (**Fig. 2n**) are consistent with those based on the full SCRS (**Fig. 2b, e**). When using only these two peaks to model single-cell starch and TAG, strong within-population starch-TAG NC was absent at the early stage of N-(0h-1d, 3d), whereas emerged at later stage of N-(2d, 4d-8d; **Fig. 2o, p**). These results, which are consistent with full-SCRS-derived findings above, raise the possibility of treating the intensity of each band as a potential phenotype and pairwise correlating all the ~1600 bands in a SCRS among cells in a ramanome.

### Genetically validating IRCA by knockout and then complementation of starch synthetic genes

To validate such IRCA-based prediction of inter-metabolite conversions, we employed a genetic approach (**Fig. S3; Fig. 3**; **Table S1**). CC4325, a mutant directly derived from CC124 by X-ray mutagenesis, is deficient of starch due to *sta1-1* knockout ^24^, thus the starch-TAG conversion (i.e., NC between starch and TAG bands in IRCA) should attenuate. As expected, IRCA of time-series CC4325 ramanomes revealed that the strong starch-TAG NC among single-cells was no longer present (0-96h under N-; **Fig. S3b**), which is in contrast to CC124 (**Fig. 2o, p**).

The cell wall of *Cr* contains cellulose whose Raman signal can overlap with that of starch, which potentially interferes with the reliability of using the 940 cm^−1^ band to quantify single-cell starch content. To test this possibility, we collected time-series ramanomes for CC406, a wall-less strain derived from CC124 ^25^. For CC406, IRCA revealed strong NC (*ρ* < −0.6) starting at 6h under N− and remained so afterwards, suggesting starch-to-TAG conversion then (**Fig. S3c**). Thus, cell wall does not interfere with the ability to detect starch-TAG conversion via IRCA.

Similarly, for CC4324 (i.e., *cw15 arg7-7*), a cell-wall deficient, arginine-requiring, starch-producing strain derived from CC406, significant NC took place throughout the 96h (especially at 24h, 72h and 96h; **Fig. 3a**). In contrast, for CC4333 (i.e., *sta6-1*, **Fig. 3b**) or CC4334 (i.e., *sta7-1*, **Fig. S3g**) which both are direct CC4324 derivatives yet produce no starch (due to disruption of *sta6* and *sta7* respectively ^25^), no strong starch-TAG correlation (*ρ* > −0.6) was reported by IRCA over the full course (0-96h under N-).

Interestingly, for CC4326 (i.e., *sta1-2*), another direct CC4324 derivative whose starch synthesis pathway is only partially disrupted ^25^, the temporal correlation pattern in IRCA is distinct from CC4333 and CC4334 which are fully starchless (under N-; **Fig. 3c**): instead of the latter’s full ablation of starch-TAG NC throughout 96h, CC4326 exhibited strong starch-TAG NC yet only at the mid phase of from 12h to 48h, consistent with transient accumulation and conversion of a low level of starch to TAG and cease of the conversion upon starch depletion. Notably, under N+ which serves as a control condition (no TAG accumulate in *Cr* under abundant medium N ^25^), CC4326 did not exhibit the starch-TAG NC until at the very late phase (e.g., 96h; when medium N was depleted by algal growth); in contrast, for CC4333 and CC4334, no starch-TAG NC is apparent throughout the 96h under N+, consistent with their genotypes (i.e., complete disruption of starch synthetic pathway; **Fig. S4**; ^25^).

Importantly, for both CC4565 and CC4566, both complemented strains with fully restored starch synthetic capability (i.e., the disrupted *sta6* was re-introduced into CC4333 by overexpression via pSL18-STA6; ^25^), the temporal patterns of starch-TAG NC under N- are identical to CC4324 (i.e., the direct progenitor of CC4333, carrying an intact starch synthetic pathway): significant NC throughout the 96h (under N-; **Fig. 3d, e**). Altogether, the starch-TAG NC in IRCA that indicates starch being converted to TAG is directly correlated with starch-synthetic genotypes. Thus IRCA can detect and model the interconversion between two metabolites from a single instance of isogenic population.

### IRCNs reveals novel product-product and substrate-product links

In each ramanome, the 1581 Raman bands in each SCRS represents over one million pairwise correlation among individual cells; such rich information indicate an enormous number of possible between-phenotype links and in particular, between-metabolite conversion. For example, for the CC124 N-7d ramanome that consists of 60 cells (which exceeds the minimal sampling depth for IRCA; **Supplemental Information**; **Fig. S5**), all pairwise among-cell correlations of bands unveiled totally 60994 strong NC ‘links’ (*ρ* < −0.6, *P* < 0.05) that formulate a 1581-node network (**Fig. 4a**; **Methods**). In such Intra-ramanome Correlation Networks (IRCN), a node represents a Raman band in SCRS and thus a potential phenotype (e.g., metabolite), while a link indicates a tentative interaction between two phenotypes (e.g., conversion).

**Figure 4.**
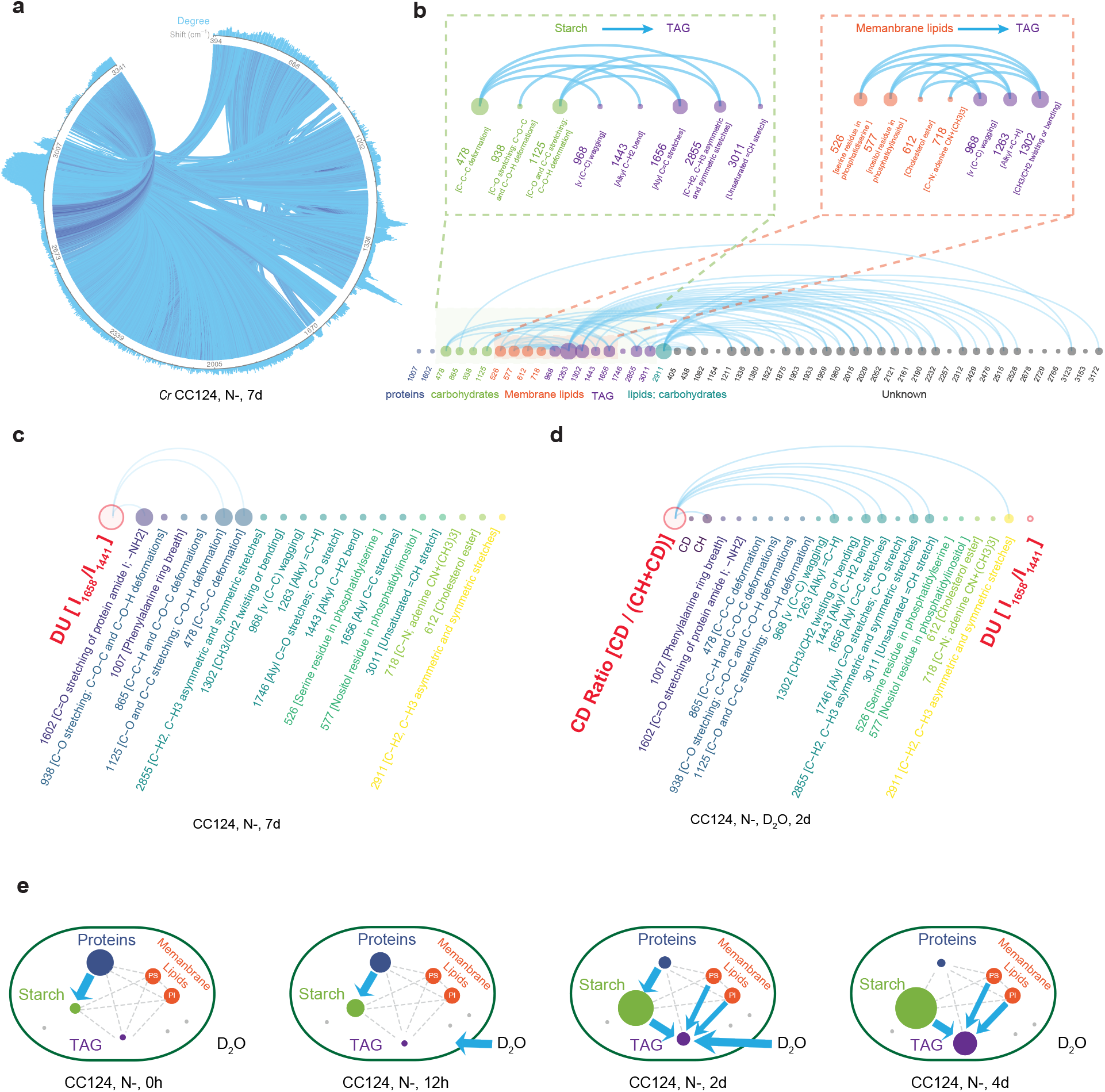
Global and local IRCN of wild-type *C. reinhardtii*. (**a**) Global-IRCN of 1581 Raman bands of CC124 under N-at Day 7 (*ρ* < −0.6, *P* < 0.05). (**b**) Local-IRCN with 51 Raman peaks of CC124 under N- at Day 7. (**c**) Local-IRCN (i.e., 17 characteristic Raman peaks) that includes DU of CC124 under N- at Day 7 (*ρ* < −0.6, *P* < 0.05). (**d**) The local-IRCN that further includes the CD ratio of CC124 under 25% D2O and N- at 2d. Edge represent strong negative correlations (*ρ* < −0.6, *P* < 0.05). N-: nitrogen depleted condition. (**e**) Key substrate-product or product-product links discovered by IRCNs. Contents of starch and TAG were quantified by 940 cm-1 and 2851 cm-1 respectively. Protein content was modeled via full SRCS. Degree of lipid unsaturation (DU) was quantified by the ratio between 1658 cm^−1^ (unsaturated C>C bonds) and 1441 cm^−1^ (saturated C–C bonds). D2O incorporation rate was quantified by the CD Ratio, i.e., the ratio between C-D bonds area (2040 to 2300 cm-1) and C-D + C-H bonds area (2040 to 2300 cm-1 and 2800 to 3050 cm-1). Blue lines and blue arrows represent strong conversions.

An IRCN is information-rich. For example, in a 51-peak module of CC124 N-7d IRCN (**Fig. 4b**; **Fig. S6a**), besides 478 cm^−1^ and 938 cm^−1^ for starch and 1658 cm^−1^ and 2853 cm^−1^ for TAG, many bands of known or unknown assignments are present, suggesting additional metabolite conversions. In particular, 526 cm^−1^ (phosphatidylserine; PS) and 577 cm^−1^ (phosphatidylinositol; PI) exhibit strong NC with 968 cm^−1^, 1302 cm^−1^ and 1263 cm^−1^ which are all TAG markers ^12^ (**Fig. 4b**). As both PS and PI are main components of membrane lipids, this observation suggests the conversion between membrane lipids and TAG at N-7 days ^26, 27^, which is supported by (*i*) the concomitant TAG accumulation and membrane lipid degradation under stress for *Cr* ^28^ and (*ii*) stable, modest rise of cellular FAME levels yet dramatic fall of chloroplast membrane lipid level at early stage of N- in *Cr* ^29, 30^. Notably, comparison of the 16 time-series IRCNs of CC124 under N-revealed increasing trends of the PS-TAG and PI-TAG correlations (**Fig. S6b-e**). Moreover, the NC between membrane lipids and TAG occurred as earlier as 8h, much earlier than that between of starch and TAG (**Fig. 2o, p; 2d**). Notably, in the N-7d IRCN, the starch-TAG NC is also present, indicating starch-TAG conversion (**Fig. 4b**). Thus the local IRCN module reveals multiple concomitant conversion processes that all produce TAG.

Notably, this network can be dynamically expand to accommodate additional nodes that represent phenotypes underpinned by multiple bands. For example, the degree of lipids unsaturation (DU), a key feature determining the application and value of lipids, can be quantified by the ratio between 1658 cm^−1^ (unsaturated C=C bonds) and 1441 cm^−1^ (saturated C–C bonds) ^12, 31, 32^. Inclusion of DU into the time-series IRCNs revealed a local module of 17 peaks with functional assignment, where DU is highly negatively correlated with two starch bands (938 cm^−1^ and 1125 cm^−1^) and one protein band (1007 cm^−1^) at the late phase of N-(**Fig. 4c**; **Fig. S7a, b**; **2d-8d**). This suggests starch and proteins are converted to unsaturation lipids at the late phase of N-, consistent with the concordant (*i*) degradation of proteins and starch and (*ii*) accumulation of unsaturation lipids ^14, 19, 20, 21, 22, 23^.

Moreover, IRCN can reveal interactions between substrate intake and products. Intake of D_2_O by cell resulted in substitution of C-H bonds in macromolecules in the cell by C-D bonds; as a result, CD ratio, i.e., the ratio between C-D bond area (2040 to 2300 cm^−1^) and area of C-D plus C-H bonds (2040 to 2300 cm^−1^ and 2800 to 3050 cm^−1^), can measure cellular metabolic activity ^33, 34^. In a separate time-resolved experiment, D_2_O was fed to CC124 immediately after removing medium nitrogen (i.e., ramanome sampled in triplicates at 0h, 12h, 1d, 2d, 3d and 4d under N-; **Fig. S6c**; **Methods**). IRCA revealed, (*i*) specifically at 2d, a module of 17 characteristic peaks was formulated in which CD ratio exhibit strong NC (*ρ* < −0.6, *P* < 0.05) with five TAG bands (1263, 1443,1656, 2855 and 3011 cm^−1^; **Fig. 4d, Fig. S6c, d**), suggesting that at 2d higher-TAG-content cells exhibit lower D_2_O-assimilating activity (and vis versa). Thus 2d represents a most ‘diverse’ metabolic state where both low-TAG, active-‘drinking’ cells and high-TAG, inactive-‘drinking’ cells co-exist, while before 2d the former dominates, and after 2d the latter dominates. (*ii*) Between CD ratio and DU, no strong NC CD (*ρ* < −0.6, *P* < 0.05) was observed at any timepoints (**Fig. 4d, Fig. S6c, d**), except for weak correlation at 3d (*ρ* = −0.50, *P* < 0.01), suggesting DU of synthesized lipids increased with reducing *Cr* vitality under N- stress (supported by GC-MS data ^12^). (*iii*) Bands of starch (the major carbon storage form of *Cr* under N-) showed no correlation with CD ratio at any timepoints. Thus starch synthesis appears mainly supported by endogenously derived H, while TAG synthesis require exogenously supplied H. Therefore, IRCA is a new strategy to track cellular destination of target substrate.

Altogether, a choreography of interplay among water-intake and major cellular products was revealed (**Fig. 4e**): (*i*) at 0h, only protein-starch conversion took place; (*ii*) at 12h, both protein-starch conversion and D_2_O incorporation occurred; (*iii*) at 2d, besides protein-starch conversion, TAG became the most catabolically active components, and starch, PS, PI and D_2_O all contributed to TAG synthesis; (*iv*) at 4d, the three conversions of starch-TAG, PS-TAG and PI-TAG still occurred, but not protein-starch and D_2_O-TAG. Therefore, IRCA unveils metabolic dynamics from a single snapshot of an isogenic population.

### Global features of IRCNs reveal the degree of dynamics for the metabolite-conversions

To probe the global features of IRCN, key network parameters were derived for the time-series IRCNs of CC124 under N-(*ρ* < −0.6, *P* < 0.05; **Fig. 5**; heatmap of *ρ* shown in **Fig. S8a**). (*i*) The number of nodes (*num_Node*; **Fig. 5a**) was increasing (min of 122 at 2h and max of 1499 at 7d), so did the number of edges (*num_Edge*; *ρ* < −0.6, *P* < 0.05; **Fig. 5b**). (*ii*) The number of modules (*num_Module*; each module is a sub-IRCN not connected with any other nodes in the IRCN; **Fig. 5c**) was decreasing (max of 11 at 4h and min of 1 at 18h, 2d, 4d, 6d and 7d). However, size of the largest module (*size_largest_Module*; number of nodes in the largest module of each IRCN; **Fig. 5d**) increased greatly (min of 53 at 2h and max of 1499 at 7d). (*iii*) Both density (*Density*, number of edges divided by number of all possible edges for the same nodes; **Fig. 5e**) and average degree (*ave_Degree*, average number of adjacent edges; **Fig. 5f**), both depicting network complexity, have increased. In contrast, average *PCC* (*ave_PCC*, **Fig. 5g**) turned more negative (from −0.000078 at 2h to −0.0335 at 7d), indicating more frequent and active conversions between the metabolites implicated.

**Figure 5.**
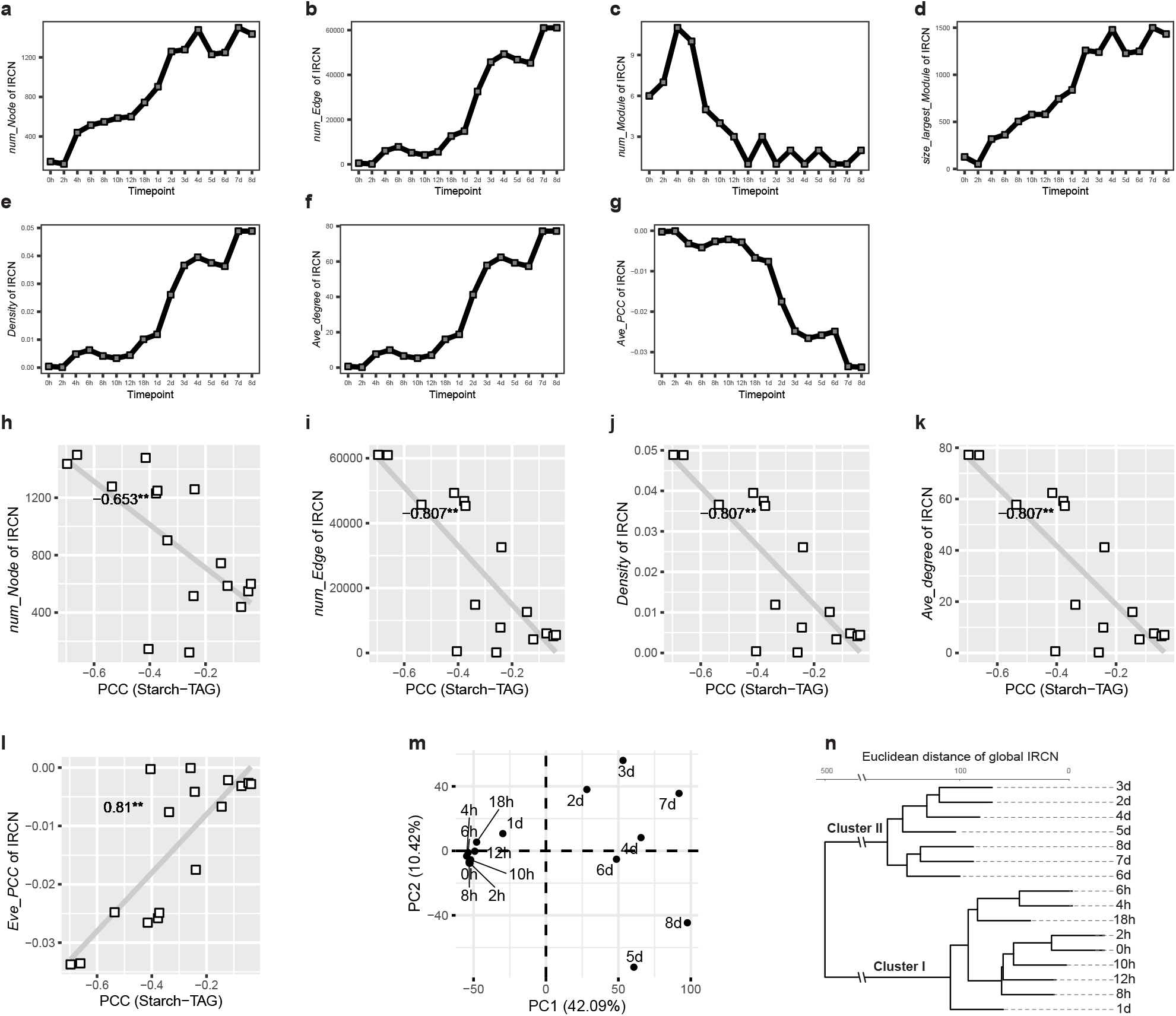
Intra-Ramanome Correlation Network (IRCN) reveals global feature of metabolite conversion dynamics. (**a-g**) Key network parameters derived from the time-series IRCNs (*ρ* < −0.6, *P* < 0.05) of CC124 under N-, including number of nodes (*num_Node*, **a**), number of edges (*num_Edge*, **b**), number of modules (*num_Module*; each module is a sub-IRCN not connected with any other nodes in the IRCN, **c**), size of the largest module (*size_largest_Module*; number of nodes in the largest module of each IRCN, **d**), density (*Density*, ratio of the number of edges divided by the number of all possible edges of the same nodes, **e**) and average degree (*ave_Degree*, average number of adjacent edges, **f**), average PCC (*ave_PCC*, sum of all significant strong negative correlation divided by all nodes, **g**). (**h-l**) Correlations between starch-TAG conversion mode (PCC of starch-TAG) and key network parameters, i.e., *num_Node* (**h**), *num_Edge* (**i**), *Density* (**j**), *ave_Degree* (**k**), and *ave_PCC* (**l**). Also shown are clusters of IRCNs via PCA (**m**) and HCA (**n**) based on the IRCN correlation matrix (strong negative correlations *ρ* < −0.6, *P* < 0.05).

Across all time points, dynamics of *num_Node*, *num_Edge*, *Density*, *ave_Degree* and *ave_PCC* of 16 IRCNs were significantly correlated with starch-TAG conversions (*PCC* of −0.653, −0.807, −0.807, −0.807 and 0.81 respectively, *P* < 0.05, **Fig. 5h-l**). In addition, Raman bands at the carbohydrate and lipid region were usually prominent among the nodes with highest degree (**Fig. S9**). Moreover, the 1457 cm^−1^ region (the alkyl C–H_2_ bend of saturated lipids) dominated at the later phases under N-(i.e., 4d, 5d, 7d and 8d), consistent with the turnover between lipid classes (saturated switching to unsaturated). Therefore, topological features of an IRCN can reveal both metabolically active compounds and their inter-conversions.

Based on the matrix of *ρ* (*ρ* < −0.6, *P* < 0.05), the 16 CC124 IRCNs under N- can be classified as Clusters I and II (**Fig. 5m, 6n**): I consists of time points before 24h, which are characterized by global features of low *num_Edge* and low intensity of *ave_PCC et. al*; II includes those after 24h, with global features of a sharp increase of *num_Edge* and *ave_PCC*. Thus 24h is the transition point (consistent with population-level measurement; **Fig. S1**), when protein stopped degradation, while starch stopped accumulation and the early build-up of starch began to contribute to other cell activities and serve as carbon source for lipid synthesis.

**Figure 6.**
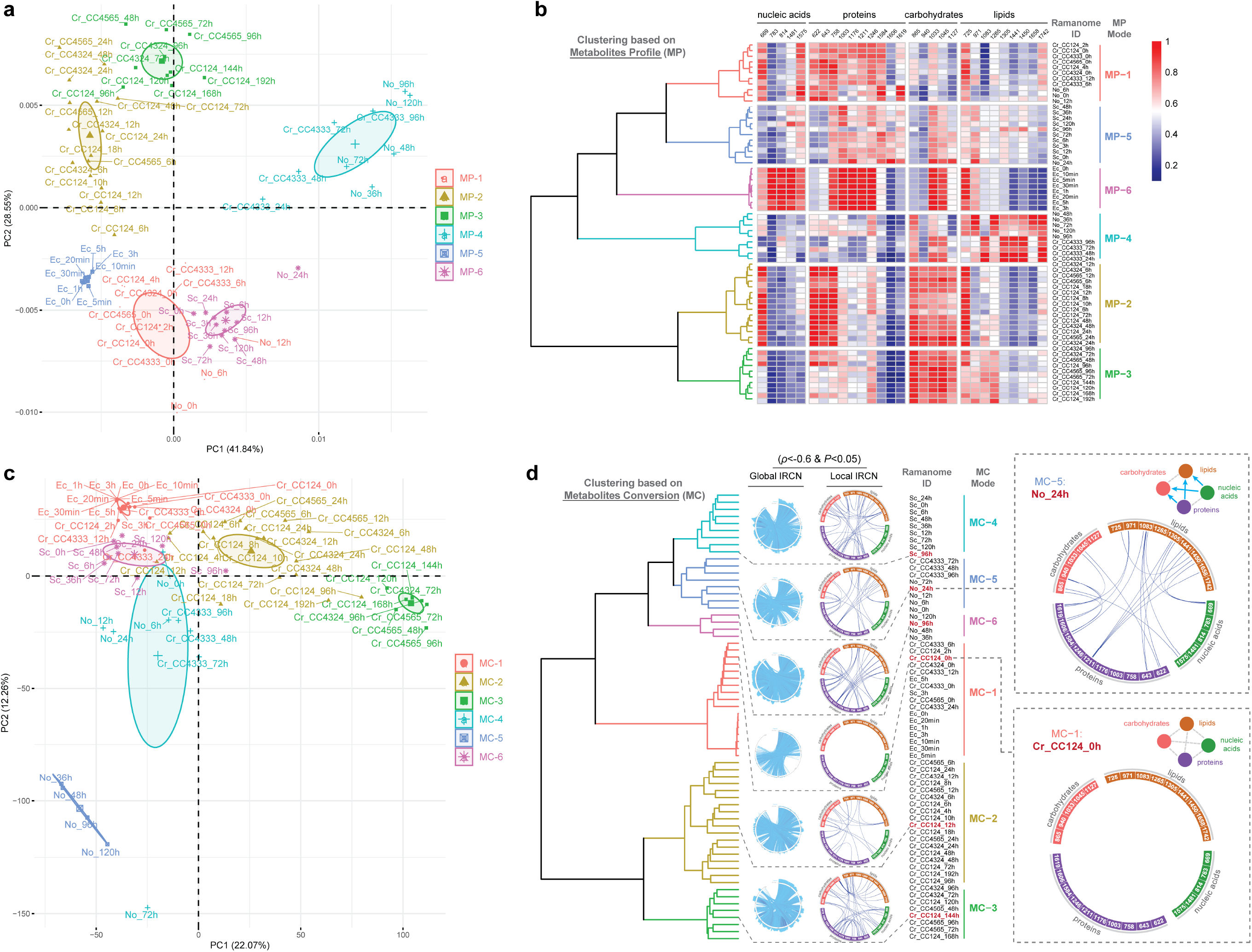
Clustering of ramanomes via the metabolite profiling (MP) and metabolite conversion (MC) signatures. PCA and HCA clusters of 64 ramanomes based on **MP** signature (mean SCRS, **a, b**) and **MC** signature (*ρ* < −0.6, *P* < 0.05, **c, d**) were shown. Details for each of the populations of *Chlamydomonas reinhardtii* (WT and starchless mutant series, *Cr*), *Nannochloropsis oceanica* (*No*), *Saccharomyces cerevisiae* (*Sc*) or *Escherichia coli* (*Ec*) are provided in **Table S1**. Details for the Raman barcodes and local IRCNs are provided in **Table S2**. Global IRCNs consist of Raman bands from 600 cm-1 to 1800 cm-1. Clusters are colored based on HCA.

In addition, the global heatmap of Δ*ρ* (the difference between max *ρ* and min *ρ* in a ramanome series) reveals the dynamic change of the “hotspots” of metabolite conversion. For example, for the 16-timepoint CC124 N-process (**Fig. S8b**), the Δ*ρ* between 1441 cm^−1^ (alkyl C-H_2_ bend, CH_2_ scissoring and CH_3_ bending; i.e., saturated lipids) and 1402 cm^−1^ (bending modes of methyl groups; i.e., proteins) is one of the most prominent hotspots (1.18; highlighted in **Fig. S8b**), with max *ρ* of 0.619 at 2h (strong positive correlation) and min *ρ* of −0.557 at 7d (strong NC). This suggests the temporal switch, from synergic degradation of lipids and protein at 2h to protein-to-lipid conversion at 7d, is one key defining dynamic features of this process.

Furthermore, comparison among such heatmaps of *ρ* can measure the global similarity of metabolite conversion modes among ramanomes from different strains (**Fig. S8c**). For example, at N-72h, IRCNs of the aforementioned CC4324 (wild-type), CC4333 (starchless mutant), CC4565 (complement strain) reveal much higher similarity with CC4324 for CC4565 than for CC4333, with the distinction of CC4333 being the absence of regions (dotted rectangle) found in CC4324 and CC4565, such as (*i*) 867-970 cm^−1^ (starch converting to lipids), and (*ii*) 1441-1658 cm^−1^ and 1441-1744 cm^−1^ (both saturated lipids converting to unsaturated lipids; solid rectangles in **Fig. S8c**). These IRCN-derived phenotypes are consistent with strain genotypes (**Fig. 3**).

### IRCN is a new “state”-specific metabolic signature of isogenic population for diverse microbes

To maximally extract metabolic features from a ramanome (**Fig. 1**), three signatures are proposed: metabolite profile (**MP**; via the mean SCRS of a ramanome; **Fig. 6a, b**), metabolite interaction (**MI**; via correlation matrix of a ramanome with significant pairwise correlations of all Raman bands (*P* < 0.05); **Supplemental Information; Fig. S10a, b**) and metabolite conversion (**MC**; via correlation matrix with strong NC of pairwise Raman bands (*ρ* < −0.6, *P* < 0.05); **Fig. 6c, d**). To test their generality, **MP, MI** and **MC** were derived for each of 64 ramanomes from *Cr* (WT and the starchless mutant series), *Nannochloropsis oceanica* (*No*), *Saccharomyces cerevisiae* (*Sc*), *Escherichia coli* (*Ec*) (**Table S1**; **Methods**).

Similarity-based clustering of **MP** revealed six modes (**Fig. 6a, b**). (*i*) **MP-1** consists of PST-0h, 2h and 4h, plus early-phase under N- of *Cr* and *No*, indicative of high protein content but low content of starch or TAG. (*ii*) **MP-2** includes PST-6h, 8h, 10h, 12h, 18h, 24h, 48h and 72h (i.e., starch-producing *Cr* strains (CC124, CC4324 and CC4565) at the middle phases (i.e., 6h-48h) under N-), indicating high carbohydrates content. (*iii*) **MP-3** includes PST-96h, 120h, 144h, 168h, 192h (i.e., mainly starch-producing *Cr* strains (CC124, CC4324 and CC4565) at the later phase (i.e., 48h-192h) of under N-), indicating high carbohydrate content and the start of accumulating lipids (mainly neutral lipids, e.g., TAG). (*iv*) **MP-4** includes both middle and later phases of *No* under N-(36h, 48h, 72h, 96h, 120h) and later phase of starchless *Cr* strains (CC4333) under N-(24h, 48h, 72h, 96h): corresponding to high content of neutral lipids and low levels of nucleic acids, protein and carbohydrates. (*v*) *Sc* (0h-120h) form **MP-5:** high protein yet low carbohydrates and lipid amounts. (*vi*) *Ec* under kanamycin (0h-5h) form **MP-6,** featuring higher amounts of proteins and nucleic acid. **MC** also formulate six modes (**Fig. 6c, d**). (*i*) **MC-1** : The PST 0h, 2h-ramanomes (mainly consisting of early phase of *Cr* strains under N− and all *Ec* states): exhibiting no or few pairwise strong negative correlation happened in these IRCNs. (*ii*) **MC-2** : PST-4h, 6h, 8h, 10h, 12h, 18h, 24h. 48h,72h and 96h (plus starch-producing *Cr* strains at the early or middle stages under N-): protein is converted into starch (**Fig. S1a**). **MC-2** showed more NC links between Raman peaks with functional assignment than **MC-1**, indicating more active metabolites-metabolites conversion. (*iii*) **MC-3**: PST-120h, 144h and 168h (i.e., the late stages of starch-producing *Cr* strains under N-when starch is converted into lipids; **Fig. 2b**), showing more and stronger NC links in IRCN than **MC-2**. (*iv*) **MC-4** includes all the *Sc*, which showed different nucleic acids-proteins-carbohydrates-lipids (NPCL) conversion patterns. (*v*) Mode **MC-5** mainly consists of later phase of the starchless CC4333 and early phase of *No* strains under N-, indicating a similar metabolite-conversion pattern of lipid-production. (*vi*) **MC-6** mainly consists of later phase of *No* strains in N-where protein is converted into lipids.

The **MP/MI/MC** signature, although inherent linked, can be highly distinct (**Fig. S11**). **MP** appear to show more species-specificity, while **MI** and **MC** more state-specificity. For example, *Ec* are clustered with algae and yeast in **MC-1**; in contrast, *Ec* are solely clustered as **MP-6**. On the other hand, **MC** can be more sensitive in detecting cellular state change than **MP** and **MI**. For example, at 4h under N-, the metabolite content of CC124 has yet to be changed (i.e., clustered with 0h in **MP-1**; it did not switch to **MP-2** until at 6h), yet the mode of metabolite-metabolite conversion has already altered (i.e., clustered with 6h-96h; **MC-2**).

## Discussions

Exploiting a fundamental and inherent property of all cellular systems, i.e., intercellular metabolic heterogeneity, here we proposed and biochemically and genetically validated the IRCA approach, which unveils a comprehensive, landscape-like network of potential interactions among metabolic phenotypes from a single instance of a living population, rather than requiring a time- or condition-series of samples. This ability is of profound implications for designing phenotyping experiment, as spatiotemporal or condition-resolved sampling can frequently become a constraint.

Moreover, from an IRCN, numerous hypotheses of inter-metabolite conversions can be generated simultaneously, based on pairwise inter-cellular correlation of Raman bands or Raman-band-derived phenotypes. The scope of such phenotypes is broad ^11, 35^, including substrate intake ^36^, product synthesis ^12, 37^ and response to environmental stress (e.g., antimicrobial susceptibility; ^10, 34^). Based on an updated list of such SCRS-modeled phenotypes, IRCNs can be constructed and interpreted to mine the ramanome data space for new inter-phenotype links without *a priori* hypotheses. Therefore, IRCA presents a new dimension of “metabolic phenome” (and highly specific signature of metabolic activity) for a cellular system that captures not just the profile of metabolites but their dynamic interactions, which enables a data-driven research strategy for profiling cellular metabolism.

The strength of IRCA also lies in its high throughput, low consumable cost, excellent scalability and general applicability. As SCRS do not require labeling cells with chemical or genetic probes, all cells live or dead can be tackled. Moreover, acquisition of SCRS can be automated (e.g., via flow-mode Raman cytometry or sorting; ^38, 39^), thus IRCA can be readily extended to the plethora of cell types under diverse culture conditions, including historical ramanome data.

On the other hand, the value of IRCN is dependent on breadth and accuracy of models that associate one or a combination of Raman band to a metabolic phenotype. Although the list of such models is rapidly growing ^11^, at present, only a small portion of Raman peaks in a SCRS is assigned to a metabolic phenotype. Moreover, as a result, specific biological assignment for many of the edges (i.e., those phenotype pairs showing significant NC) in the IRCN remains difficult, as bands for different compounds can overlap. Therefore, to establish new assignments or improve their specificity, new experimental and computational methods, such as stable-isotope probed SCRS (to track target-substrate assimilation) and multivariate curve resolution-alternating least squares (MCR) algorithm (to deconvolute macromolecular components from overlapping Raman bands ^40, 41^), should be developed. Nevertheless, its high throughput, low cost, excellent scalability and general applicability suggest broad use of IRCA for metabolic interrogation of cellular systems.

## Materials and Methods

### Strains and growth conditions

For *Chlamydomonas reinhardtii*, a total of 9 wild-type (WT), low-starch mutant and starch-less mutant strains were employed (**Table S1**). Specifically, CC124, CC406 and CC4324 were WT strains. The low-starch mutant CC4325 was derived by X-ray mutagenesis from CC124. The low-starch mutant CC4326 and the starch-less mutants CC4333 and CC4334 were derived from CC4324 by random integration of cassette pARG7 in the nuclear genome ^42, 43, 44^. The starch-less mutant CC4333 is deficient in the catalytic (small) subunit of ADP-glucose pyrophosphorylase, which interrupts synthesis of the ADP-glucose, a substrate for starch biosynthesis. The CC4334 mutant contains a disrupted isoamylase gene thus starch level is severely attenuated, but it accumulates a soluble glycogen-like product. CC4565 and CC4566 were complemental strains of CC4333, derived by complementation with plasmid pSL18-STA6. All these mutants can be obtained from the Chlamydomonas Resource Center (http://www.chlamycollection.org).

The *C. reinhardtii* cells were inoculated into TAP (Tris acetate phosphate) liquid medium with or without arginine (100 μg mL^−1^) supplement under one-side continuous light (approximate 150 µmol photons m^−2^ s^−1^) at 25°C bubbled with air to ensure mixture and to prevent settling and grown to late log phase in N-replete TAP medium. Then they were re-inoculated at 1×10 ^6^ cells/ml in parallel into nitrogen-replete TAP medium (N+), nitrogen-depleted TAP medium (N-; in which NH_4_Cl was omitted), or 25% D_2_O nitrogen-deplete TAP medium (25% D_2_O N-), each in triplicates. Cultures at each of a series of timepoints were sampled, in triplicates (**Table S1**).

For the *Nannochloropsis oceanica* IMET1 strain, cells were cultured in a modified f/2 liquid medium with 4 mM NO_3_^−^ under continuous light (approximately 50 μmol photons m^−2^ s^−1^) at 25°C and induced in nitrogen-replete (N+) or nitrogen-depletion (N-) f/2 medium; the cultures were sampled at multiple timepoints. For *E. coli* DH5α, cells were cultured in a glass tube with fresh LB medium (10 g/L NaCl, 10 g/L Tryptone, 5 g/L Yeast extract, 3.7 μg/mL kanamycin) at 37°C in a shaking incubator (130 rpm); samples were collected at 0, 5, 10, 20, 30, 60, 180 and 300 min ^10^. For *Saccharomyces cerevisiae* Y50049, cells were cultured in a glass tube of fresh YPD medium at 30°C in a shaking incubator (200rpm); samples were collected at 0h, 3h, 6h, 12h, 24h, 36h, 48h, 72h, 96h, 120h. All cultures and all sampling for further analysis were in triplicates.

### Acquisition of Single-cell Raman spectra from isogenic population of cells

Raman spectra of individual cells were acquired using a modified Raman Microscpectroscopy equipped with a confocal microscope with a 50 ×PL magnifying dry objective (NA=0.55, BX41, Olympus, UK) and a 532 nm Nd:YAG laser (Ventus, Laser Quantum Ltd, UK). The scattered photons were collected by a Newton EMCCD (Andor, UK) utilizing a 1600 × 200 array of 16 µm pixels with thermoelectric cooling down to −70°C for negligible dark current. Before measurement, each sample was washed three times and resuspended in ddH_2_O to remove the culture media. For algal and yeast samples, cells were loaded in a capillary tube (50mm length× 1mm width × 0.1mm height, Camlab, UK). The power out of the objective was 100 mW. For each SCRS, the signal acquisition time was 2 seconds for *C. reinhardtii* and *N. oceanica*, and 3 seconds for yeast. For *C. reinhardtii* and *N. oceanica*, prior to Raman signal acquisition, the cell was quenched with 532 nm laser until the signal of fluorescent and resonantly enhanced biomolecules was no longer detectable. For each individual cell, a background spectrum was generated as the average of four spectra acquired from the liquid around the cell. For *E. coli* samples, cells were loaded onto a clean CaF_2_ slide and air dried before Raman measurement, and the SCRS *E. coli* samples were collected as described ^10^.

### Intra-Ramanome Correlation Analysis

The raw SCRS was first pre-processed with LabSpec 5 (Horiba Scientific, France), including background subtraction and baseline correction by a polynomial algorithm (with degree of seven). Then SCRS were normalized, followed by IRCA which used a computational pipeline for data analysis and result visualization.

Due to potential technical variation (e.g., change of data collector, batch, etc), SCRS from different ramanomes may have distinct spectral ranges and resolution. Therefore, prior to computing the MP signature and the MI and MC networks, the 64 ramanomes were standardized in three steps: (*i*) For spectral range, only the "fingerprint area" (600-1800 cm^−1^) was extracted; (*ii*) spectral resolution was simulated to 1 cm^−1^, via the interpolation algorithm; (*iii*) spectral normalization was performed by division by its area [10, 12]. The temporal ramanome dataset of CC124 under N-along 16 timepoints was regarded as a standard Protein-Starch-TAG (PST) metabolism process, which was then "projected" to the entire 64 ramanomes dataset.

For each IRCN, the correlation matrix of each ramanome was constructed by calculating the Pearson Correlation Coefficient (*PCC*; *ρ*) of all possible pairwise combinations of Raman bands, among all the 60 cells sampled for the ramanome. Those pairs of Raman bands with significant correlation was used as candidates that potentially indicate metabolite interactions (*P* < 0.05), while those with strong negative correlations suggesting potential conversion among two metabolites (*ρ* < −0.6, *P* < 0.05). The *igraph* package in R (http://www.r-project.org) was used to derive the key network properties and visualize the IRCNs (either from specific subsets of Raman peaks or of all Raman bands). To probe the global features of an IRCN, key network parameters including number of nodes (*num_Node*), number of edges (*num_Edge*), number of modules (*num_Module*; each module is a sub-IRCN not connected with any other nodes in the IRCN), size of the largest module (*size_largest_Module*; number of nodes in the largest module of each IRCN), density (*Density*, ratio of the number of edges divided by the number of all possible edges of the same nodes) and average degree (*ave_Degree*, average number of adjacent edges), average PCC (*ave_PCC*, sum of all significant strong negative correlation divided by all nodes), were derived. For global IRCNs, all Raman bands were used. For simplified versions of IRCN that facilitate visualization, characteristic marker Raman peaks were used (669, 783, 814 and 1481 cm^−1^ for nucleic acids, 622, 643, 758, 1003, 1176, 1211, 1246, 1584, 1606 and 1619 cm^−1^ for proteins, 865, 940, 1033, 1045 and 1127 cm^−1^ for carbohydrates, 725, 971, 1083, 1265, 1305, 1441, 1450, 1658 and 1742 cm^−1^ for lipids; **Table S2**).

To characterize, compare or cluster ramanomes, three signatures of **MP, MI** and **MC** were proposed. For **MP**, the mean SCRS of each ramanome was used. The **MI** network of a ramanome was generated as follows: (*i*) a correlation matrix was constructed by calculating the *PCC* of all possible pairwise Raman bands; (*ii*) significant correlations (*P* < 0.05) were tabulated, while those with no significant difference were counted as 0 (no correlation). The **MC** network of a ramanome was generated by including only those significant, strongly negative correlations (*ρ* < −0.6, *P* < 0.05). To measure the similarity, pairwise Euclidean distances were calculated. Then hierarchical cluster analysis (HCA) was performed with Ward’s algorithm and six clusters produced ^45, 46^. Principal component analysis (PCA, the *factoextra* package in R) was used to visualize the **MP, MI** and **MC** signatures.

## Acknowledgements

This work was supported by Ministry of Science and Technology of China (2018YFA0902500), National Science Foundation (32030003, 31900074) and Natural Science Foundation of Shandong, China (ZR201807090262). J.X. and Y.H. introduced the IRCA concept and designed experiments. Y.H. and P.Z. performed experiments. Y.H., S.H. and Y.J. analyzed the data. J.X, and Y.H. wrote the manuscript.

