## Supplemental files for "Intra-Ramanome Correlation Analysis Unveils Metabolite Conversion Network from an Isogenic Cellular Population"

**Supplemental Materials and Methods**

**Supplemental Results**

**Supplemental References**

**Supplemental Tables**

**Table S1, S2**

**Supplemental Figures**

**Figure S1-S11**

### Supplemental Materials and Methods

#### Quantification of starch, protein and TAG content (population-level) for the *Chlamydomonas reinhardtii* wild-type strain CC124

Dry algal biomass was obtained via lyophilization by a vacuum freezing dryer. Starch content of dry algal biomass was quantified using an enzymatic starch assay kit (Amyloglucosidase/α-amylase Method, K-TSTA 07/11; Megazyme, Ireland). Briefly, samples of about 20 mg dry algal power were treated with 80% ethanol to remove sugars, hydrolyzed into soluble maltodextrins with thermostable α-amylase and then digested into D-glucose with amyloglucosidase. The glucose produced was treated with a reagent containing glucose oxidase, peroxidase and 4-aminoantipyrine and then quantified spectrophotometrically at 510 nm wavelength.

Total protein content of algal cultures was measured as previously reported [1]. Briefly, ~ 10 mg of lyophilized algal biomass was hydrolyzed in 200 μL lysis buffer (1 M sodium hydroxide, NaOH) and then incubated at 80°C for 10 min by water bath. Then 800 μL dd$H_2O$ was added to the hydrolysate to bring the volume to 1 mL. Cellular debris was centrifuged at 12,000 g for 30 min before the supernatant was transferred to a new tube. The extraction was repeated two more times and all the supernatant extracts were pooled together. Then total protein in the supernatant was determined by the BCA Protein Assay kit (cw0014s; CWBio, China) via the manufacturer's protocol.

To analyze the contents of total lipids and TAG in an algal cell population, GC-MS and TLC-GC-MS were performed. The procedures mainly include total lipid extraction, TLC, and profiling of FAMEs. Briefly, total lipids of ~30 mg lyophilized algal powder were extracted with 6mL chloroform:methanol (2:1, v/v) and recovered in chloroform. For TAG quantification, ~ 0.5 μg of lipid extract was loaded onto 10 × 20cm silica gel 60 (Merck KGaA, Darmstadt, Germany) TLC plates. TAG was separated, visualized and scraped from the plate, and then extracted with chloroform:methanol (2:1, v/v) from the TLC powder. The FAMEs were derived from all of the TAG extracts by acid-catalyzed transmethylation and then ~1 mg lipid extracts were analyzed on an Agilent 7890-5975C gas chromatography mass spectrometer fitted with a HP-INNOWAX 30 m × 0.25 mm × 0.25 μm column. The FAMEs were quantified using pentadecane as the internal standard and C8-C24 FAMEs mix as FAMEs standards.

**Quantifying the influence of sampling depth on Intra-Ramanome Correlation Analysis**

To assess the influence of sampling depth on IRCA, we first defined two parameters for an IRCN: (*i*) the cumulative mean of Pearson correlation coefficient (i.e., "*cumPCC*") between pairwise Raman peaks and (*ii*) the average degree of the IRCN (i.e., "*cumAveDegree*"). For a given ramanome, we randomly sampled the SCRS at a particular sampling depth which ranges from three to 60 cells. At each sampling depth, the trials were performed for 100000 times for *cumPCC* (10000 times for *cumAveDegree*). For each trial, *cumPCC* and *cumAveDegree* were calculated. The two parameters were then respectively plotted against the sampling depth, so as to quantitatively assess how the choice of sampling depth affects the accuracy and reliability in measuring these features.

For a ramanome, to determine whether the IRCA parameters of *cumPCC* and *cumAveDegree* are saturated at a certain sampling depth, we defined the "rate of *cumTrait*" (where "*cumTrait*" is either *cumPCC* or *cumAveDegree*) [2]:

$$\text{rate of } cumTrait\ (\mathrm{N}) = \frac{cumTrait_{\mathrm{N}} - cumTrait_{\mathrm{N}-1}}{cumTrait_{\mathrm{N}-1}}$$

N: Sample Depth; $cumTrait_{N}$: *cumTrait* at the sampling depth of N cells for a population; $cumTrait_{N-1}$: *cumTrait* at the sampling depth of (N-1) cells for a population. The relationship between the rate of *cumTrait* and sampling depth was thus plotted. We set a cutoff of 1% for the rate of *cumTrait* to define the "minimal sampling depth", which means, at this particular sampling depth, no more than 1% of gain in *cumTrait* is allowed by sampling one more cell.

**Supplemental Results**

### The effect of sampling depth affects Intra-Ramanome Correlation Analysis

To provide a rational basis for determining the proper sampling depth for IRCA, we tested the link between sampling depth and (*i*) the Pearson correlation coefficient (PCC) of pairwise Raman peaks and (*ii*) average degree of the IRCN (**Supplemental Methods**). The cumulative PCC of pairwise Raman peaks (*cumPCC*) and cumulative average degree of an IRCN (*cumAveDegree*) were derived from each of the *in silico* trials of sub-sets of SCRS sampled randomly from the full ramanome at a particular sampling depth. The parameter of "minimal sampling depth" is thus designated as the sampling depth at a cutoff of 1% for the *cumTrait* (either *cumPCC* or *cumAveDegree*), i.e., at this particular sampling depth, no more than 1% of increase in *cumTrait* will be gained by sampling one more cell (**Supplemental Methods**).

The minimal sampling depth varied for both Raman bands and timepoints. For PCC of Raman peaks of 938 $cm^{-1}$ (Starch-related Raman peak) and 2855 $cm^{-1}$ (TAG-related Raman peak), they vary from 4 (12h) to 10 (4h). For example, for the 60-cell collection at 7d, the "Minimal Sampling Depth" is 5 cells (**Fig. S5a, b**), which is much lower than our actual sample depth (i.e., totally 60 cells for each time point under each condition). For the calculation of the average Degree (i.e., a key parameter for IRCN) of the 60-cell collection at 7d, the "Minimal Sampling Depth" is 33 cells (**Fig. S5c, d**), which is also lower than the actual sample depth at each of the time points (i.e., 60 cells). Therefore, the sampling depth of 60 cells each ramanome satisfied both the pairwise links and IRCN.

### IRCN is a new "state"-specific metabolic signature of isogenic population for diverse microbes

The **MI** form six clusters (**Fig. S10a, b**). (*i*) **MI-1**: PST standard processes 0h-, 2h- and 4h-ramanome (i.e., early phase under N- of *Cr* and *No* strains), indicating that in the cells of these states among Mode **MI-1** proteins began to degrade and were converted to other metabolites (e.g. starch). (*ii*) **MI-2**: PST-6h, 8h, 10h, 12h, 18h, 24 (i.e., middle phase of starch-producing *Cr* strains (CC124, CC4324 and CC4565) under N-), indicating high carbohydrates-accumulation stage. (*iii*) **MI-3**:

PST-48h, 72h, 96h, 120h, 144h, 168h, 192h (i.e., later phase of starch-producing *Cr* strains (CC124, CC4324 and CC4565) under N-), indicating they have already accumulated high content of carbohydrates and began to convert carbohydrates to lipids. (*iv*) **MI-4**: both middle and later phases of *No* under N- (24h, 36h, 48h, 72h, 96h, 120h) and later phase of the starchless CC4333 under N- (24h, 48h, 72h, 96h), indicative of a state converting other metabolites (e.g., nucleic acids, protein and carbohydrates) to lipids. (*v, vi*) *Sc* (0h-120h) and *Ec* (0h-5h) form **MI-5** and **MI-6** respectively, under each species' characteristic metabolite-interacting networks.

### Supplemental Tables

**Table S1. Microalgal and microbial strains, culture conditions and sampled time points for IRCA in this study.**

| Species | Strains | Condition | Timepoint | Description |
|---|---|---|---|---|
| *Chlamydomonas reinhardtii* | CC124 | TAP, N- | 0h, 2h, 4h, 6h, 8h, 10h, 12h, 18h, 24h, 48h, 72h, 96h, 120h, 144h, 168h, 192h | Wild-type |
| | CC124 | TAP, N- | 0h, 6h, 12h, 24h, 48h, 72h, 96h | Wild-type |
| | CC4325 | TAP, N- | 0h, 6h, 12h, 24h, 48h, 72h, 96h | Low-starch mutant |
| | CC406 | TAP, N- | 0h, 6h, 12h, 24h, 48h, 72h, 96h | Wild-type, cell wall deficient |
| | CC4324 | TAP, N- | 0h, 6h, 12h, 24h, 48h, 72h, 96h | Wild-type, cell wall deficient |
| | CC4326 | TAP, N- | 0h, 6h, 12h, 24h, 48h, 72h, 96h | Low-starch mutant |
| | CC4333 | TAP, N- | 0h, 6h, 12h, 24h, 48h, 72h, 96h | Starch-less mutant |
| | CC4334 | TAP, N- | 0h, 6h, 12h, 24h, 48h, 72h, 96h | Starch-less mutant |
| | CC4565 | TAP, N- | 0h, 6h, 12h, 24h, 48h, 72h, 96h | CC4333 complemented strain |
| | CC4566 | TAP, N- | 0h, 6h, 12h, 24h, 48h, 72h, 96h | CC4333 complemented strain |
| *Nannochloropsis oceanica* | IMET1 | f/2, N- | 0h, 6h, 12h, 24h, 36h, 48h, 72h, 96h, 120h | Wild-type |
| *Saccharomyces cerevisiae* | Y50049 | YPD | 0h, 3h, 6h, 12h, 24h, 36h, 48h, 72h, 96h, 120h | Model eukaryotic microorganism |
| *Escherichia coli* | DH5a | LB, kan | 0min, 5min, 10min, 20min, 30min, 1h, 3h, 5h | Model prokaryotic microorganism |

**Table S2. Assignments of the Raman bands used for tracking the product profile in this study.**

| Raman Peaks | Assignments | Components |
|---|---|---|
| 622 | C-C twisting mode of phenylalanine | proteins |
| 643 | C-S stretching & C-C twisting of proteins | proteins |
| 669 | G, T | nucleic acids |
| 725 | characteristic for phospholipids | lipids |
| 758 | Tryptophan, δ (ring) | proteins |
| 783 | U, T, C | nucleic acids |
| 814 | C5’ -O-P-O-C3’ phosphodiester | nucleic acids |
| 865 | C-C-H and C-O-C deformations | carbohydrates |
| 940 | C-O stretching; C-O-C and C-O-H deformation; α-helix C-C backbone | carbohydrates |
| 971 | ν(C-C) wagging | lipids |
| 1003 | Phenylalanine ring breath | proteins |
| 1033 | υ(CO), υ(CC), υ(CCO) | carbohydrates |
| 1045 | C-O and C-C stretching; C-O-H deformation | carbohydrates |
| 1083 | Typical phospholipids | lipids |
| 1127 | C-O stretching | carbohydrates |
| 1176 | C-H in-plane bending mode of tyrosine, (CH) phenylalanine | proteins |
| 1211 | υ(C-C6H5), tryptophan, phenylalanine | proteins |
| 1246 | Amide III | proteins |
| 1265 | Alkyl =C-H | lipids |
| 1305 | CH3/CH2 twisting or bending mode of lipid | lipids |
| 1441 | Alkyl C-H2 bend | lipids |
| 1450 | CH2 bending and scissoring modes of phospholipids | lipids |
| 1481 | Nucleotide acid purine bases | nucleic acids |
| 1575 | Ring breathing modes in the DNA bases | nucleic acids |
| 1584 | Phenylalanine | proteins |
| 1606 | Phenylalanine, tyrosine, C=C | proteins |
| 1619 | υ(C=C), tryptophan | proteins |
| 1658 | Allyl C=C stretches | lipids |
| 1742 | Ester C=O stretches | lipids |

### Supplemental Figures

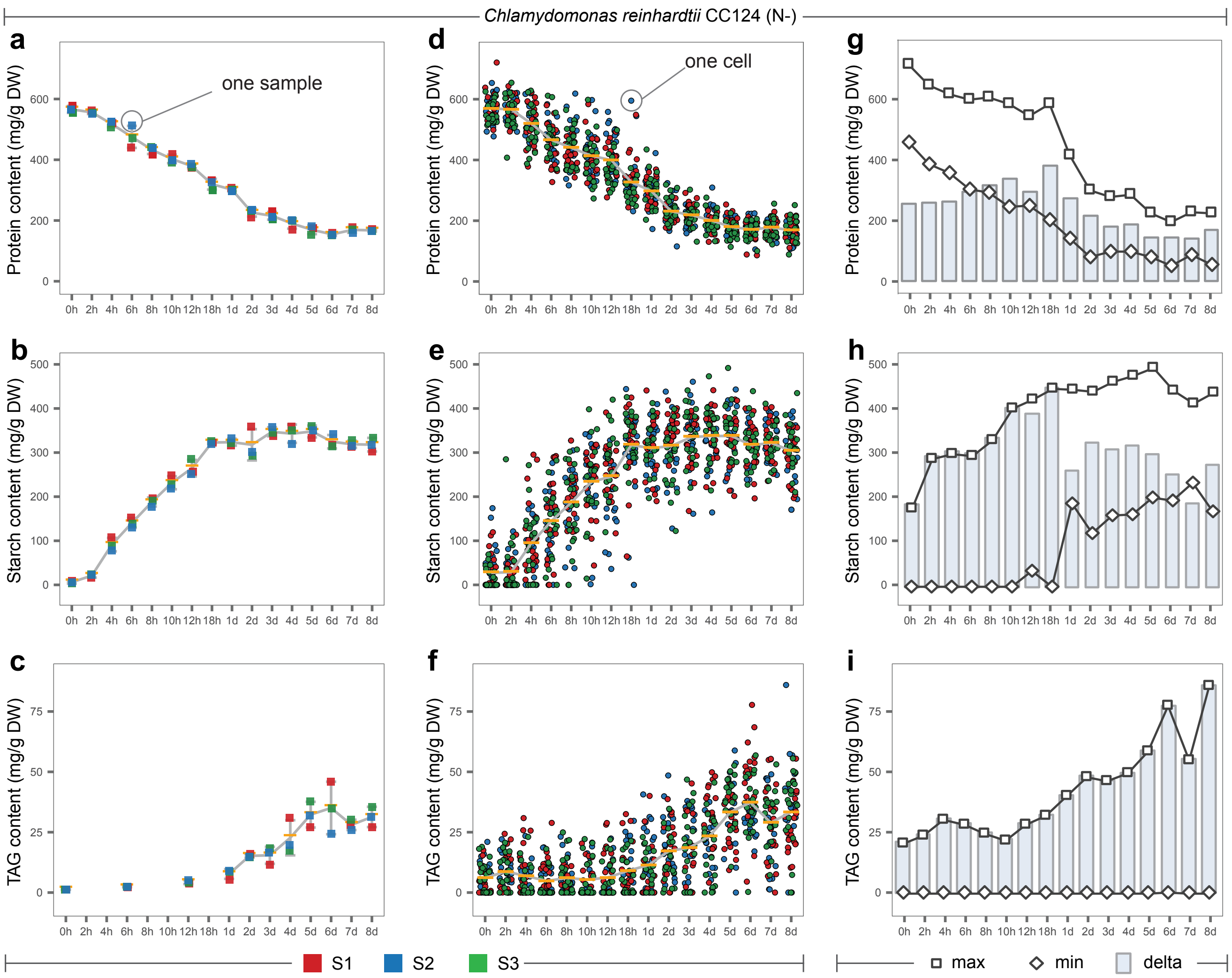


**Figure S1. Dynamics of starch, protein and TAG contents at the population and the single-cell levels.** The parameters were measured for the populations by conventional approaches (**a, b, c**) or for single cells via prediction by ramanome (**d, e, f**), max/min/delta content for each timepoints (**g, h, i**).

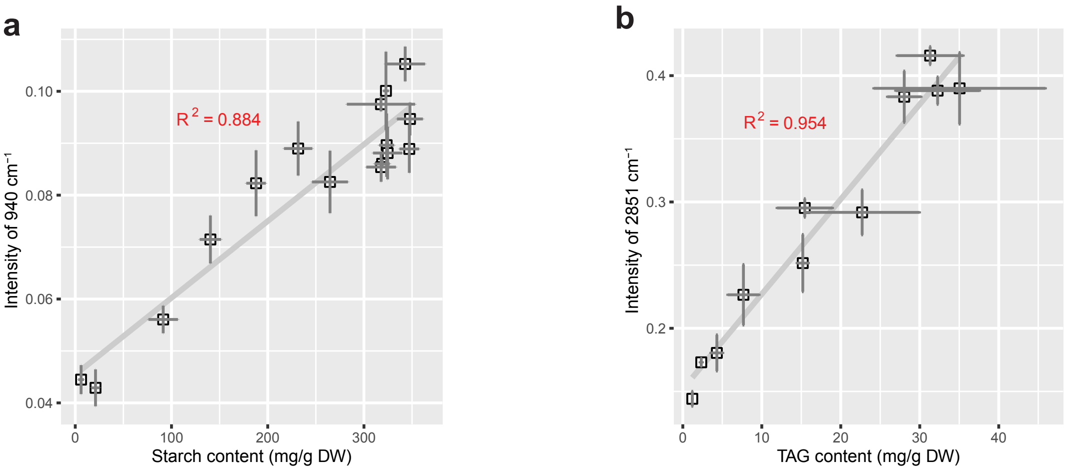


**Figure S2. Phenotypic correlation between starch and TAG contents measured by conventional methods and via intensity of 940 cm$^{-1}$ (a) and 2851 cm$^{-1}$ (b) in SCRS.**

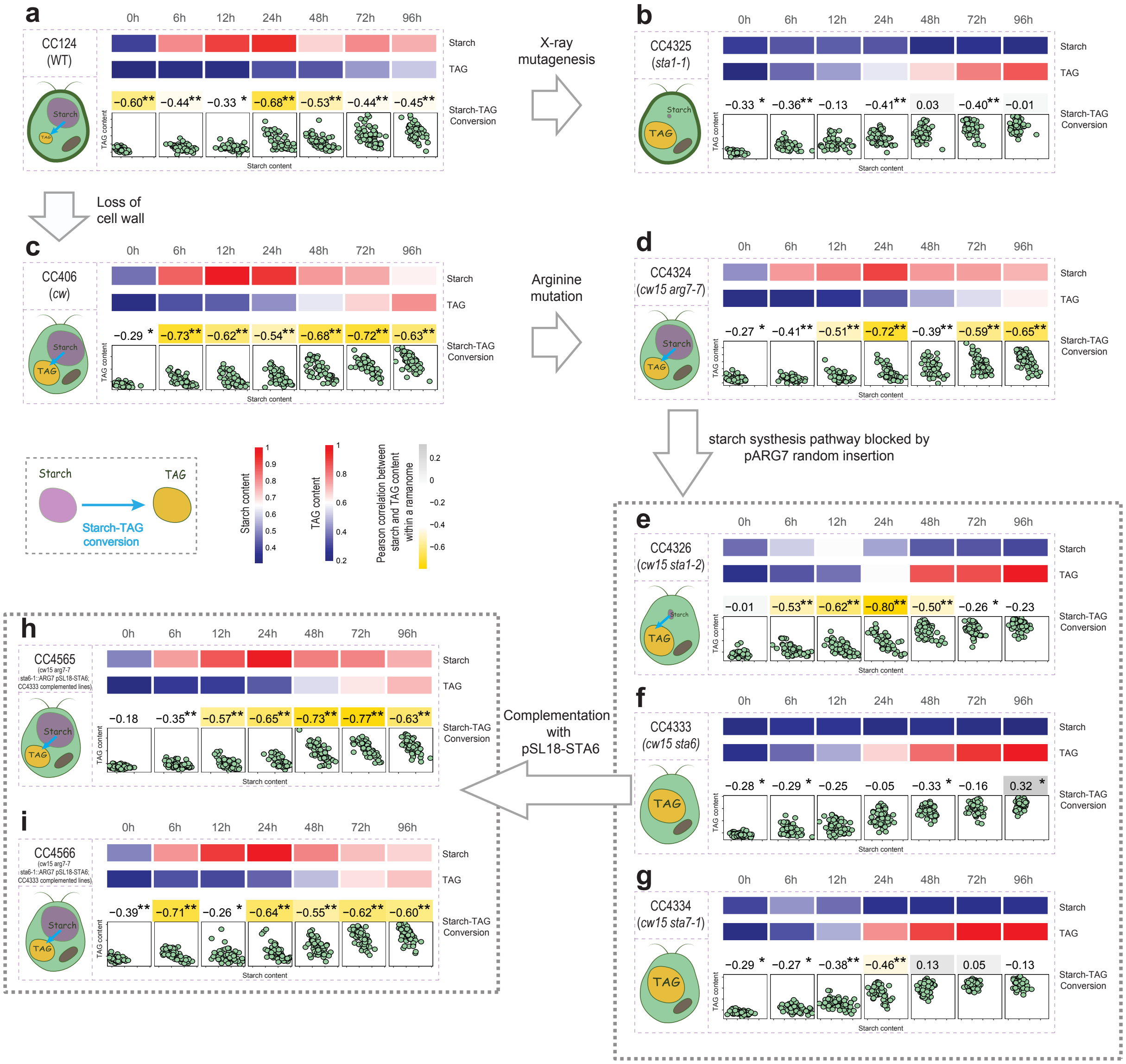


**Figure S3. Validation of the switch between starch and TAG content by multiple wild type and mutant strains of *C. reinhardtii*.** Phenotypic correlation between starch ($x$) and TAG ($y$) contents modeled by pairs of 940 $cm^{-1}$ and 2851 $cm^{-1}$ respectively for CC124 (**a**), CC4325 (**b**), CC406 (**c**), CC4324 (**d**), CC4326 (**e**), CC4333 (**f**), CC4334 (**g**), CC4565 (**h**), CC4566 (**i**) of *C. reinhardtii* strains. Detailed genotypes of the strains are listed in **Table S1**. The starch and TAG contents and Pearson correlation coefficients among the phenotypes among single cells within one ramanome were shown with heatmap (**: $P < 0.01$; *: $P < 0.05$).

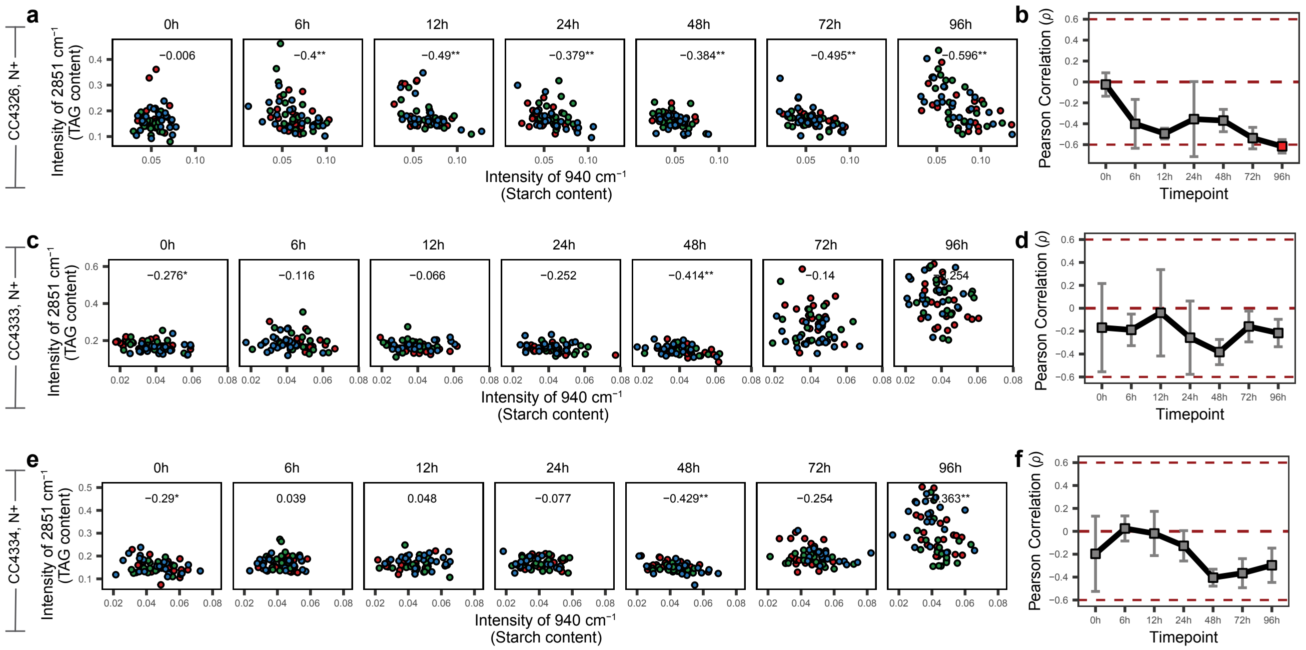


**Figure S4. Absence conversion between starch and TAG content by a directed series of mutant of *C. reinhardtii* at N+ abundant condition.** Phenotypic correlation between starch (*x*) and TAG (*y*) contents modeled by pair of Raman band of 940 $cm^{-1}$ and 2851 $cm^{-1}$ for CC4326 (**a**), CC4333 (**c**), CC4326 (**e**). In the scatterplots, each dot represents one cell. The temporal dynamics curves, *ρ* is Pearson correlation coefficients among the phenotypes among single cells (**: $P < 0.01$; *: $P < 0.05$), with value indicating mean of triplicates and error bar standard deviation for CC4326 (**b**), CC4333 (**d**), CC4334 (**f**). Absence of correlation ($\rho = 0$) or presence of strong correlation (*ρ* higher than 0.6 or lower than -0.6) was highlighted by red horizontal lines. Detailed genotypes of the strains are listed in **Table S1**.

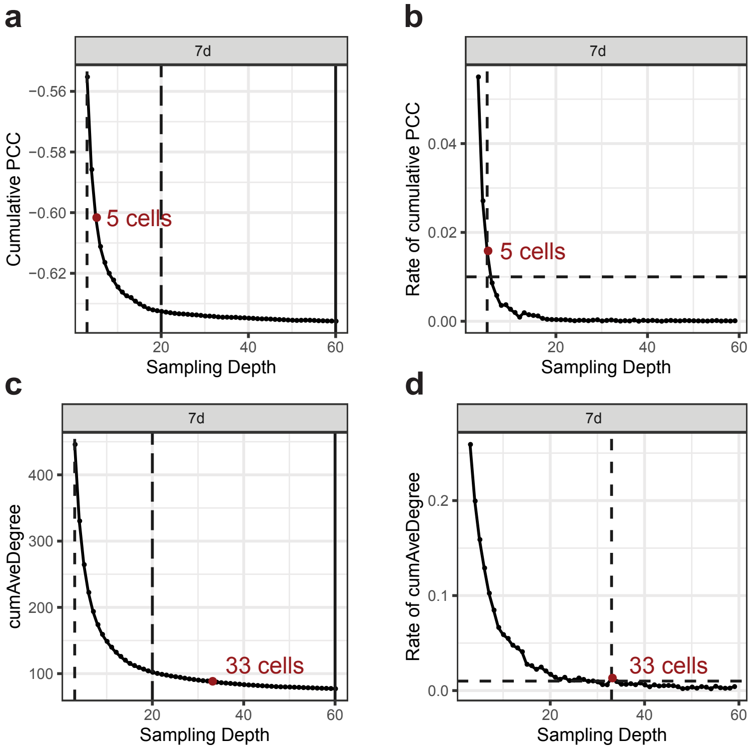


**Figure S5. The effect of sampling depth on IRCA.** Quantitative relationship is shown between sampling depth and the observed cumulative IRCA traits (*cumPCC* and *cumAveDegree*) for an *in silico* population for 60 cells in 7d-ramanome. At each sampling depth, 100000 permutations for *cumPCC* (938 $cm^{-1}$ and 2855 $cm^{-1}$) and 10000 permutations for *cumAveDegree* of sampling trials were performed, and the mean and increase rate of *cumPCC* (**a, b**) and *cumAveDegree* (**c, d**) were calculated. The minimal sampling depth, defined as the depth when no more than 1% of gain in cumulative IRCA traits is gained by sampling one more cell, was highlighted. Minimal sampling depth was high-lighted with red point and corresponding text.

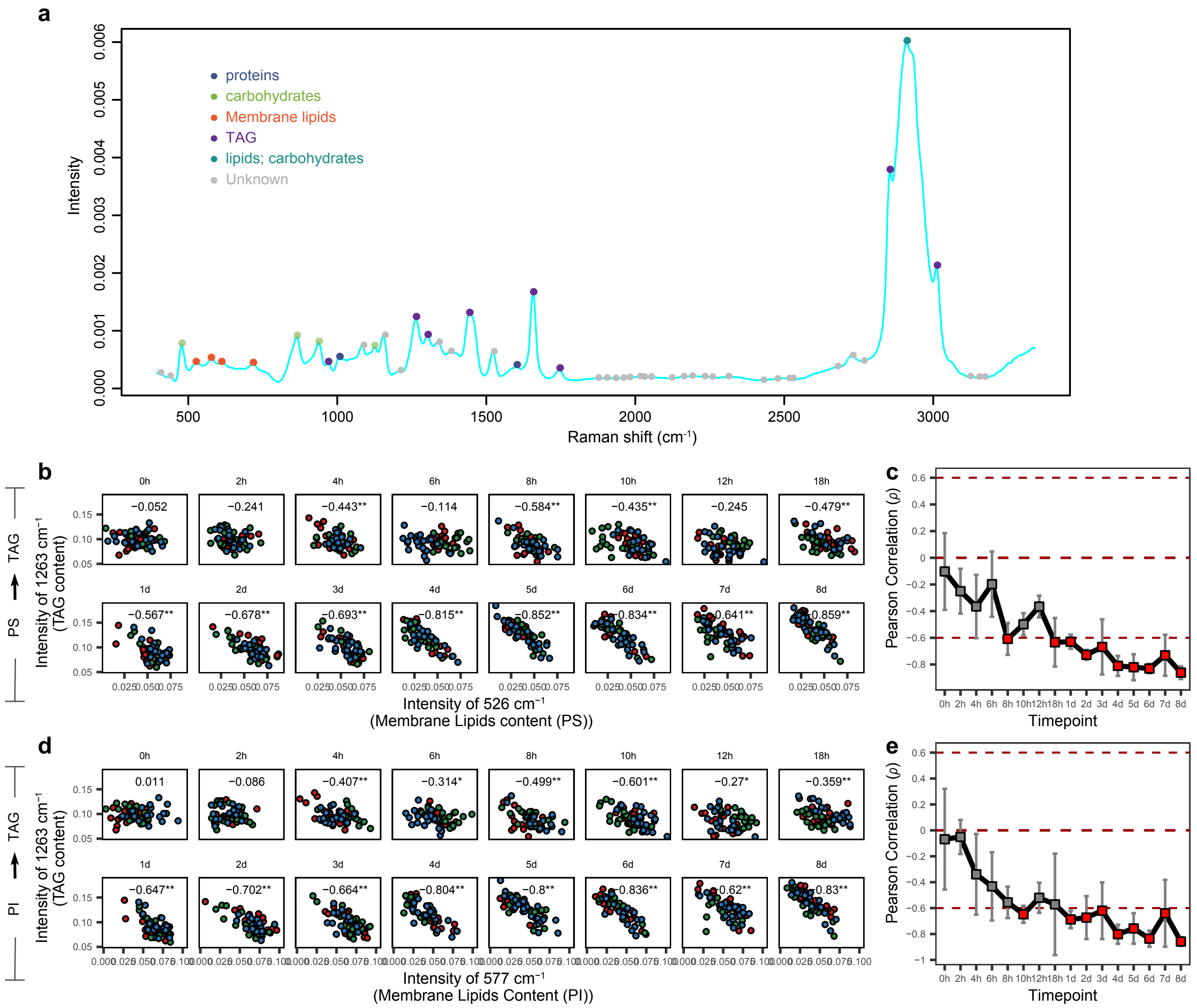


**Figure S6. Correlation of membrane-lipid and TAG contents among individual cells at each time point.** (**a**) The mean SCRS of CC124 7d ramanome (as an example), showing the major peaks and their assignments. Phenotypic correlation between membrane-lipid (*x*) and TAG (*y*) contents modeled by 526 cm$^{-1}$ (phosphatidylserine or PS) and 1263 cm$^{-1}$ (TAG) was shown (**b**, **c**), so was that for 577 cm$^{-1}$ (phosphatidylinositol or PI) and 1263 cm$^{-1}$ (**d**, **e**). In the scatterplots, each dot represents one cell. $\rho$ is Pearson correlation coefficients among single cells (**: $P < 0.01$; *: $P < 0.05$), with value indicating mean of triplicates and error bar standard deviation. Absence of correlation ($\rho = 0$) or presence of strong correlation ($\rho$ higher than 0.6 or lower than -0.6) was highlighted by red horizontal lines. Both PS and PI are main components of membrane lipids.

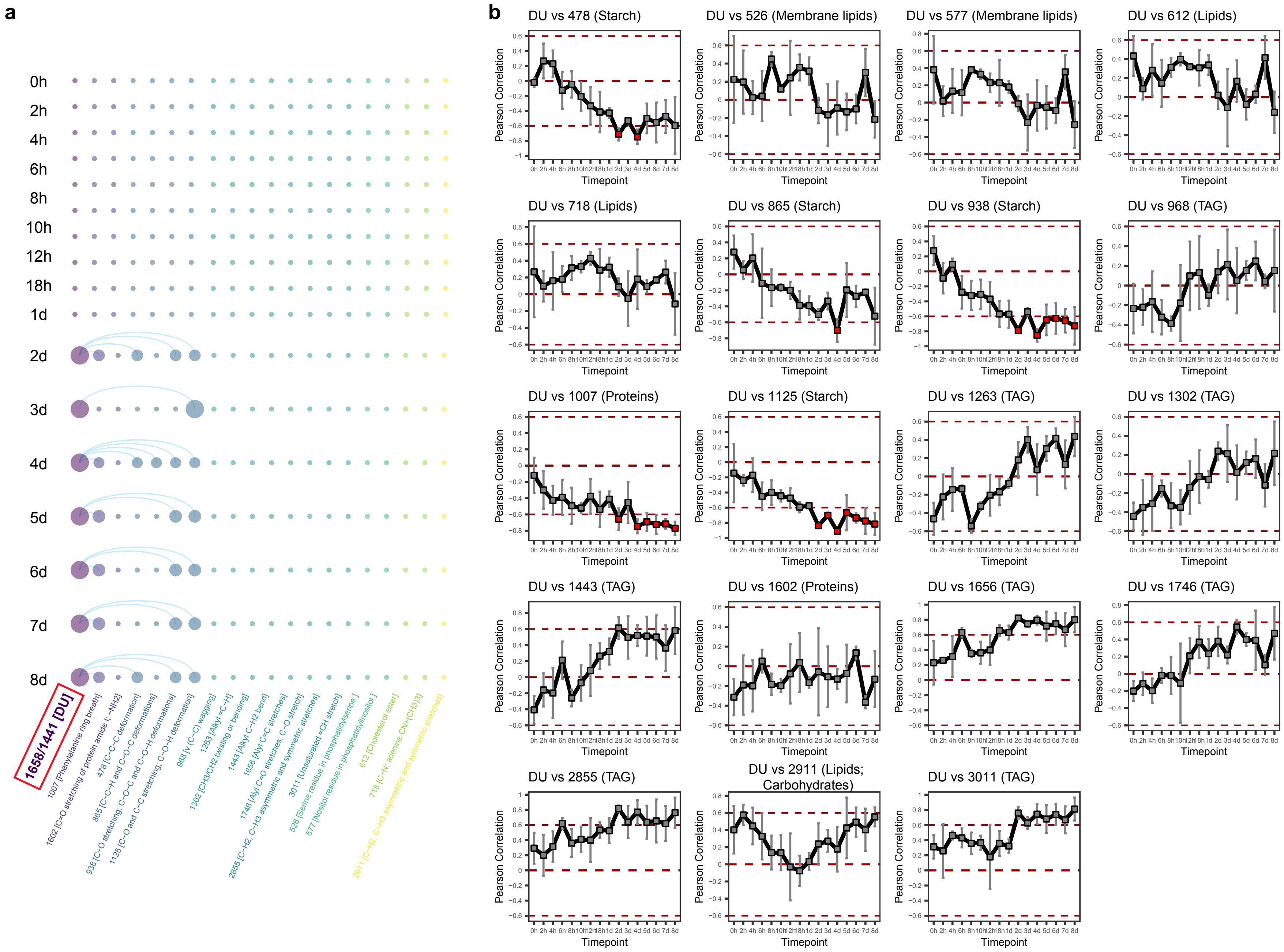
a
b
0h
2h
4h
6h
8h
10h
12h
18h
1d
2d
3d
4d
5d
6d
7d
8d
1658/1441 [DU]
DU vs 478 (Starch)
DU vs 526 (Membrane lipids)
DU vs 577 (Membrane lipids)
DU vs 612 (Lipids)
DU vs 718 (Lipids)
DU vs 865 (Starch)
DU vs 938 (Starch)
DU vs 968 (TAG)
DU vs 1007 (Proteins)
DU vs 1125 (Starch)
DU vs 1263 (TAG)
DU vs 1302 (TAG)
DU vs 1443 (TAG)
DU vs 1602 (Proteins)
DU vs 1656 (TAG)
DU vs 1746 (TAG)
DU vs 2855 (TAG)
DU vs 2911 (Lipids; Carbohydrates)
DU vs 3011 (TAG)
Pearson Correlation
Timepoint

158

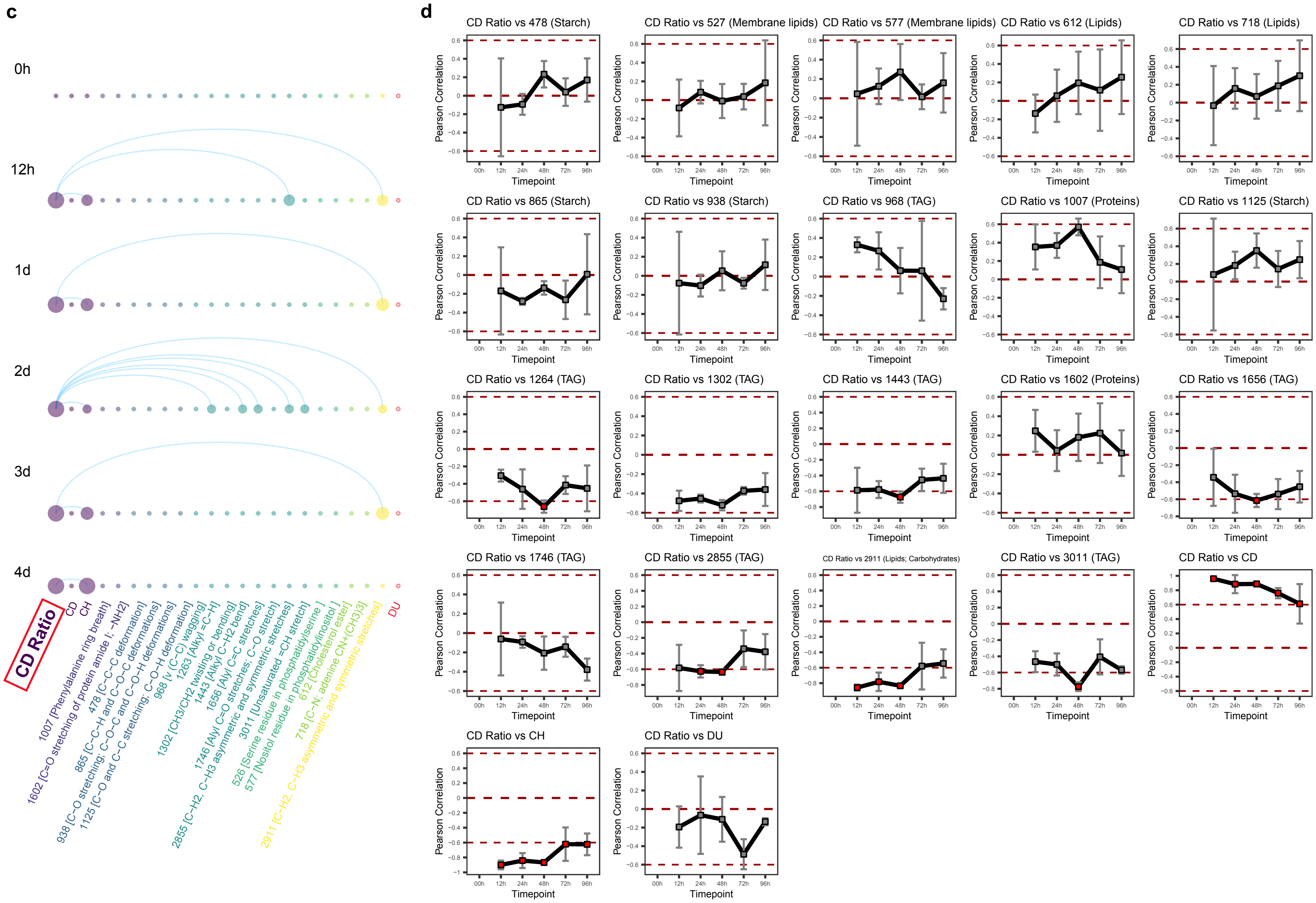
c
0h
12h
1d
2d
3d
4d
CD Ratio
CD
CH
1007 [Phenylalanine ring breath]
1602 [C=O stretching of protein amide I; −NH2]
478 [C−C−C deformation]
865 [C−C−H and C−O−C deformations]
938 [C−O stretching; C−O−C and C−O−H deformations]
1125 [C−O and C−C stretching; C−O−H deformation]
968 [ν (C−C) wagging]
1263 [Alkyl =C−H]
1302 [CH3/CH2 twisting or bending]
1443 [Alkyl C−H2 bend]
1656 [Alyl C=C stretches]
1746 [Alyl C=O stretches; C−O stretch]
2855 [C−H2, C−H3 asymmetric and symmetric stretches]
3011 [Unsaturated =CH stretch]
526 [Serine residue in phosphatidylserine]
577 [Nositol residue in phosphatidylinositol]
612 [Cholesterol ester]
718 [C−N; adenine CN+(CH3)3]
2911 [C−H2, C−H3 asymmetric and symmetric stretches]
DU
d
CD Ratio vs 478 (Starch)
CD Ratio vs 527 (Membrane lipids)
CD Ratio vs 577 (Membrane lipids)
CD Ratio vs 612 (Lipids)
CD Ratio vs 718 (Lipids)
CD Ratio vs 865 (Starch)
CD Ratio vs 938 (Starch)
CD Ratio vs 968 (TAG)
CD Ratio vs 1007 (Proteins)
CD Ratio vs 1125 (Starch)
CD Ratio vs 1264 (TAG)
CD Ratio vs 1302 (TAG)
CD Ratio vs 1443 (TAG)
CD Ratio vs 1602 (Proteins)
CD Ratio vs 1656 (TAG)
CD Ratio vs 1746 (TAG)
CD Ratio vs 2855 (TAG)
CD Ratio vs 2911 (Lipids; Carbohydrates)
CD Ratio vs 3011 (TAG)
CD Ratio vs CD
CD Ratio vs CH
CD Ratio vs DU
Pearson Correlation
Timepoint
00h 12h 24h 48h 72h 96h

**Figure S7. Local-IRCN that includes degree of lipids unsaturation (DU) and CD Ratio.** (**a**) Local-IRCN that includes the DU of CC124 at 16 time points (threshold: strong negative correlation $\rho < -0.6$, $P < 0.05$). The temporal dynamics curves of correlations between DU and 17 characteristic Raman peaks (**b**). (**c**) Local-IRCN that includes the CD ratio of CC124 under 25% $D_2O$ at 6 time points ($\rho < -0.6$, $P < 0.05$). The temporal dynamics of correlations between CD ratio and 17 characteristic Raman peaks (plus DU). (**d**). $\rho$ is Pearson correlation coefficient among the phenotypes among single cells, with value indicating mean of triplicates and error bar standard deviation. Absence of correlation ($\rho = 0$) or presence of strong correlation ($\rho$ higher than 0.6 or lower than -0.6) was highlighted by red horizontal lines.

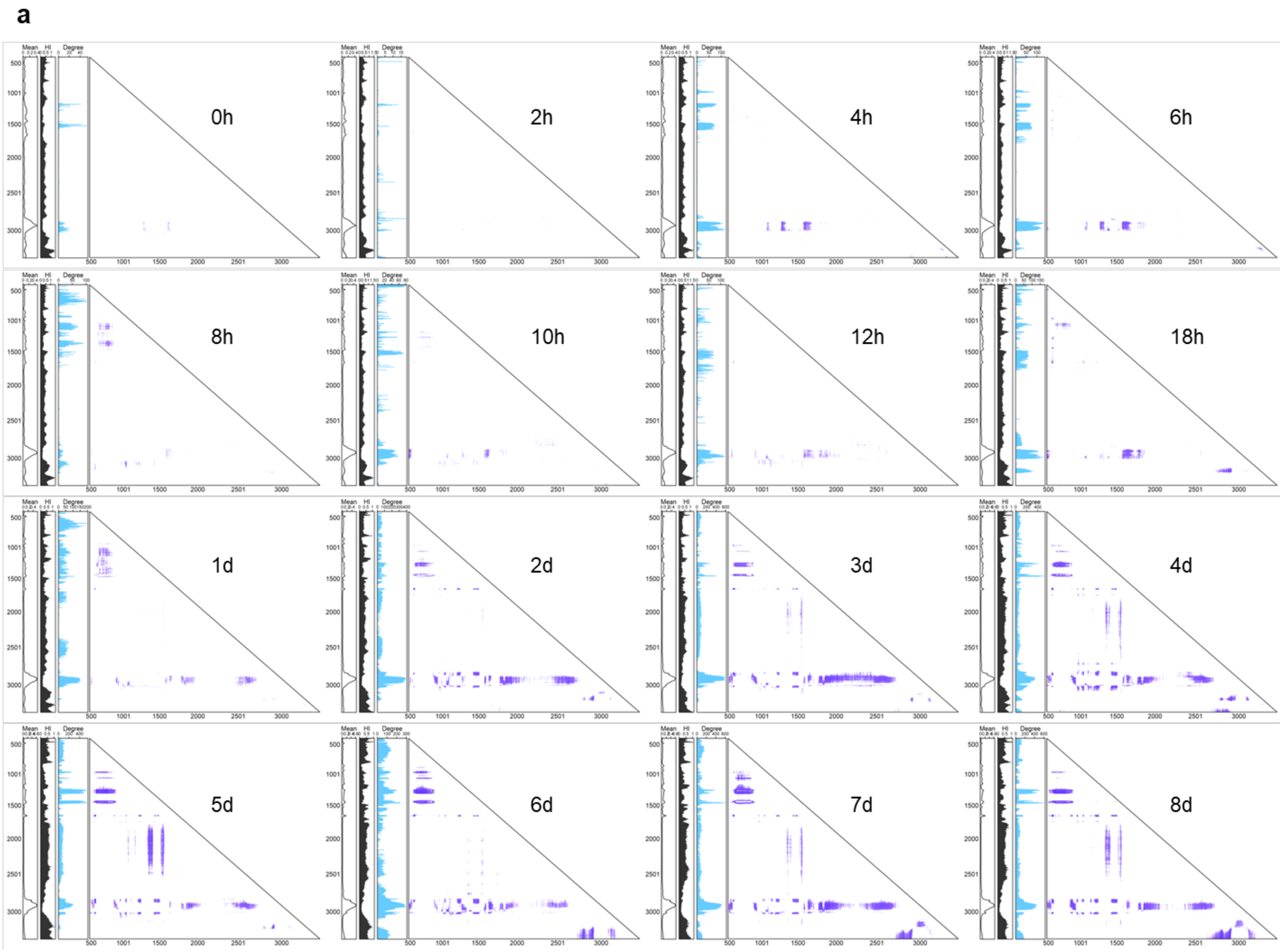
a
0h
2h
4h
6h
8h
10h
12h
18h
1d
2d
3d
4d
5d
6d
7d
8d
Mean
HI
Degree

**b**

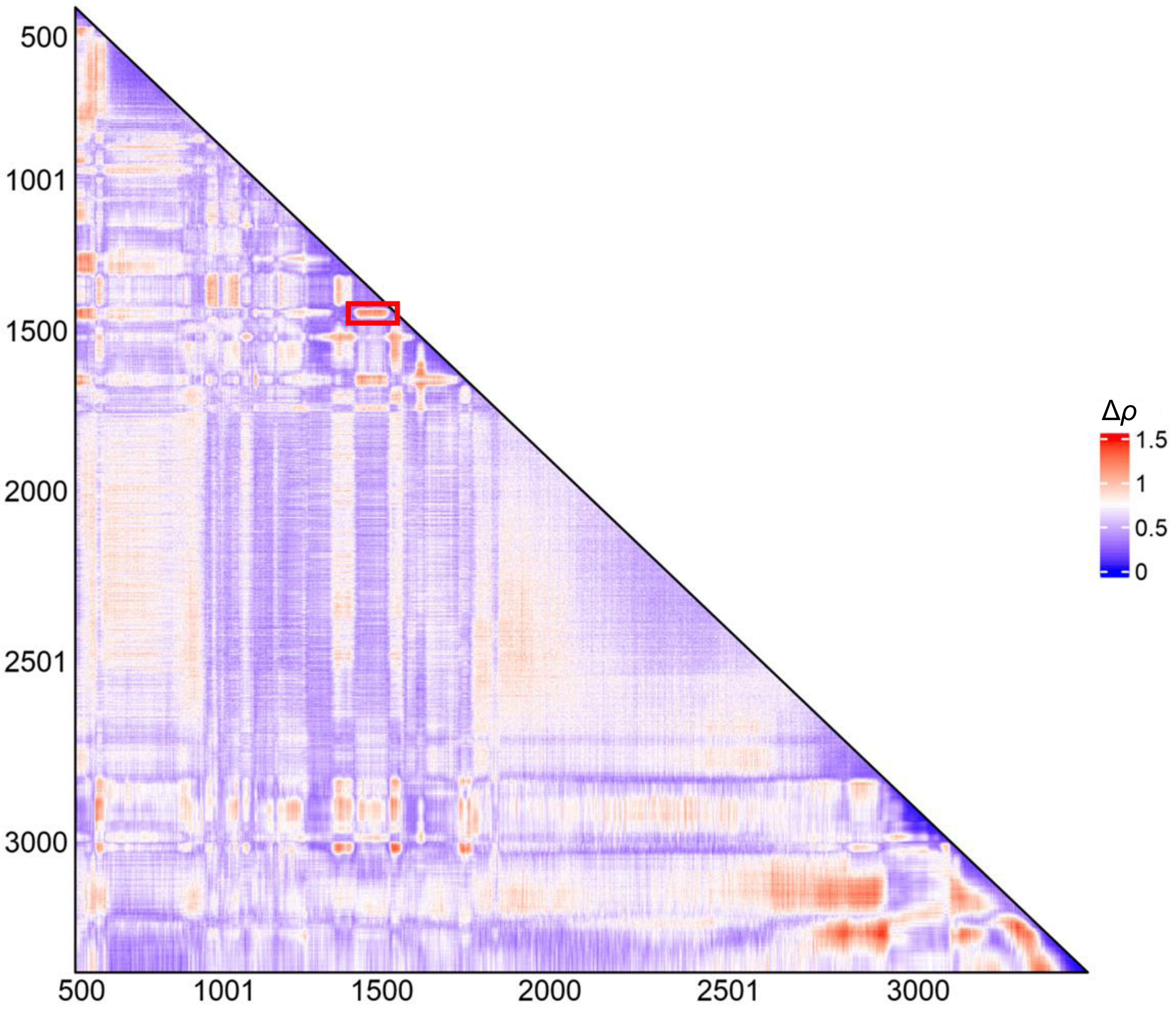

**c**

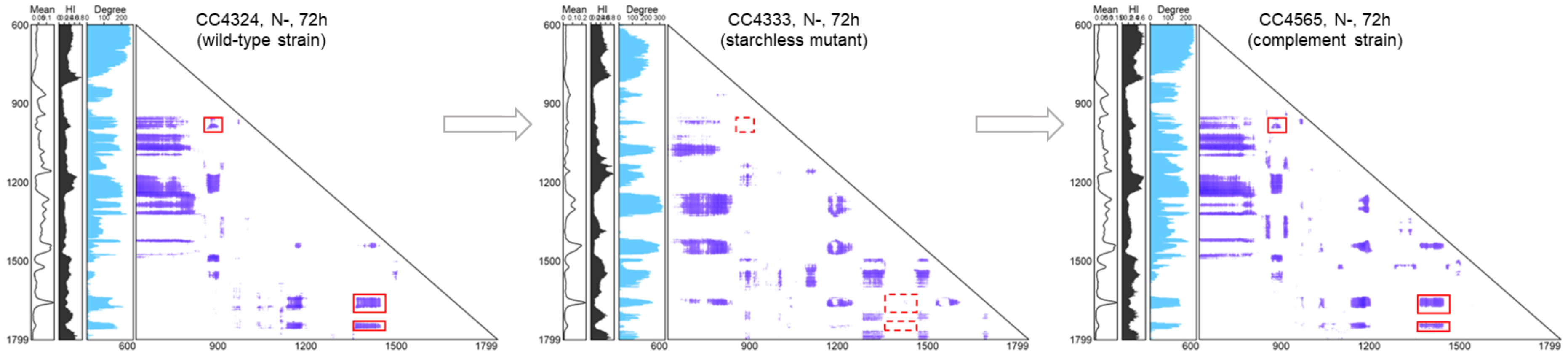


**Figure S8. Time-series IRCNs of CC124 under N- reveals global feature of metabolite conversion dynamics.** (**a**) IRCNs ($\rho < -0.6$, $P < 0.05$) were shown as 2D heatmaps. (**b**) $\Delta\rho$ of the IRCNs (i.e., 0h-8d). $\Delta\rho = \rho_{max} - \rho_{min}$. (**c**) IRCNs of three ramanomes (CC4324_N-_72h, CC4333_N-_72h and CC4565_N-_72h) in 2D heatmap give a global view of metabolite conversion modes of different species. Mean: mean spectra of one ramanome; HI: The Heterogeneity Index of a Raman band is defined as the RSD (relative standard deviation) of individual cells within a ramanome; Degree: number of degree for each bands in an IRCN.

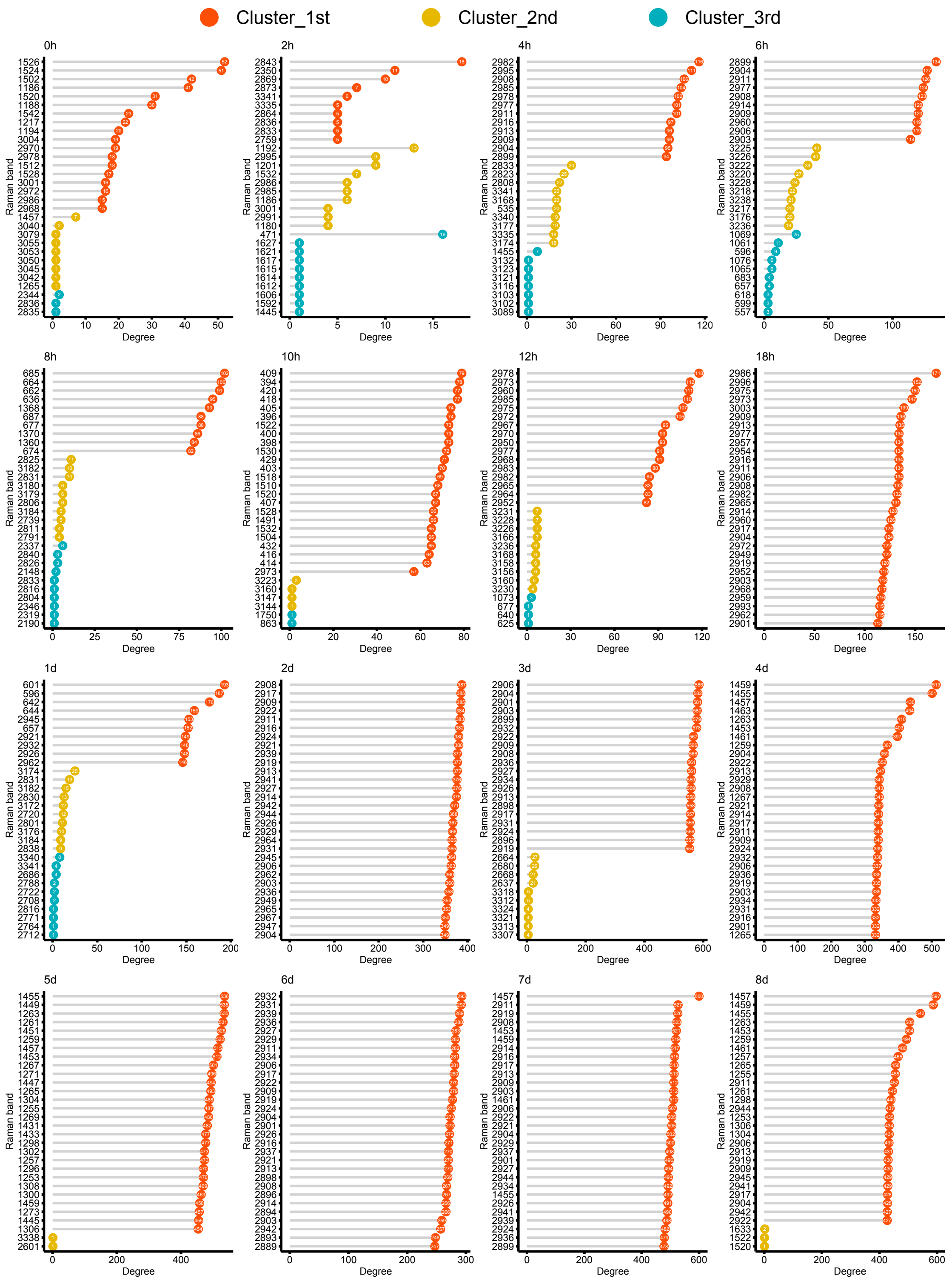


**Figure S9. Degree of the most active Raman bands for the three largest modules in each IRCN.** IRCNs were constructed with the following threshold: strong negative correlation, $\rho < -0.6$ and $P < 0.05$.

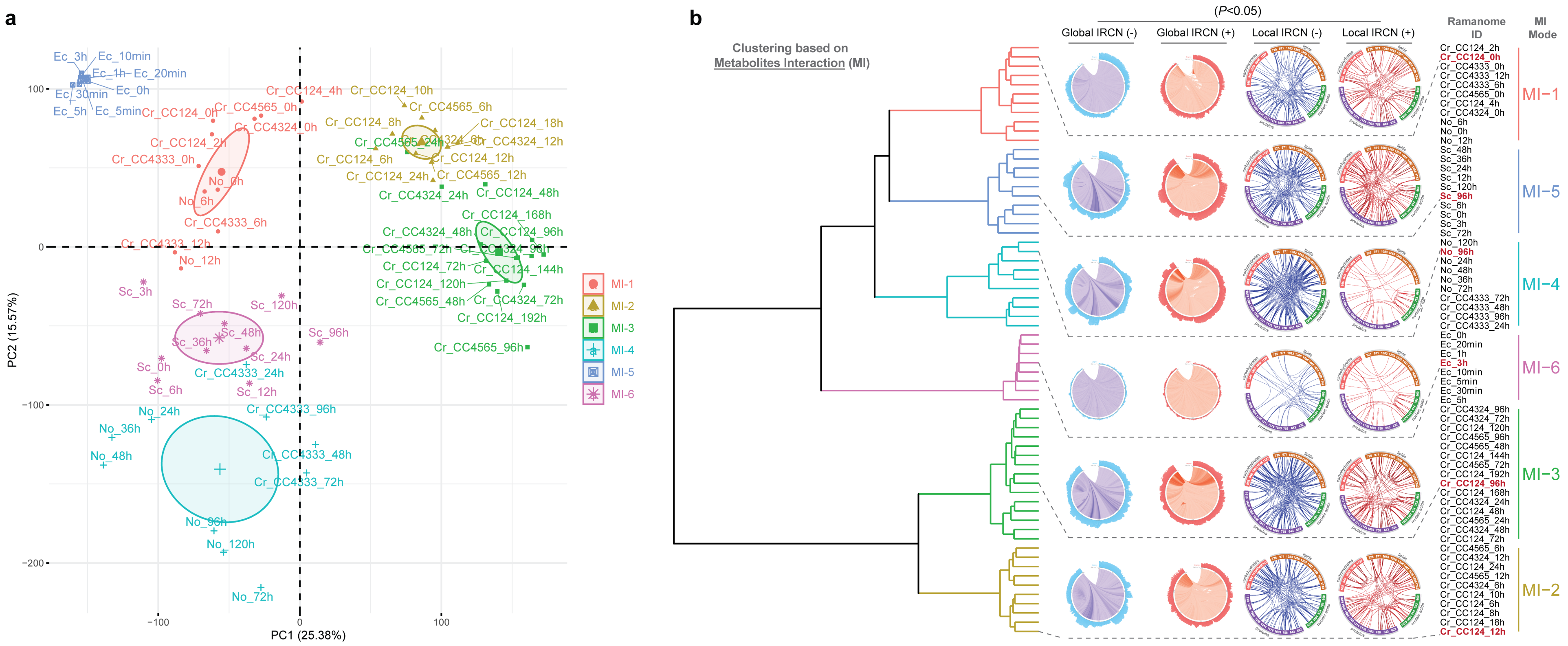


**Figure S10. Clustering of ramanomes via the metabolite interaction (MI) signatures.** PCA and HCA clusters of 64 ramanomes based on **MI** signature ($P < 0.05$; **a**, **b**) were shown. Details for each of the populations from *Chlamydomonas reinhardtii* (WT and starchless mutant series, *Cr*), *Nannochloropsis oceanica* (*No*), *Saccharomyces cerevisiae* (*Sc*) or *Escherichia coli* (*Ec*) are provided in **Table S1**. Details for Raman features of Raman barcode and local IRCNs are provided in **Table S2**. Global IRCNs consist of Raman bands from 600 $cm^{-1}$ to 1800 $cm^{-1}$. Clusters are colored based on HCA.

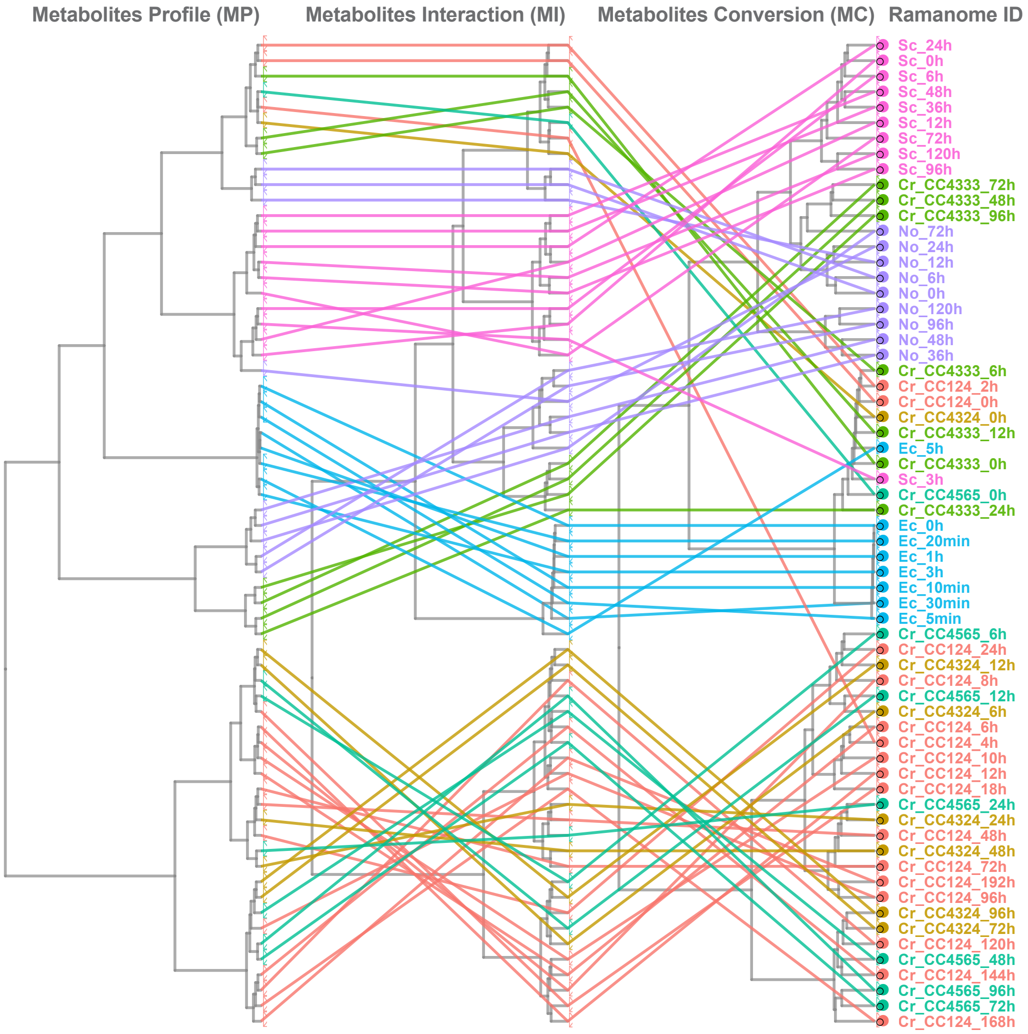


**Figure S11. Similarity and distinction among the clustering based on MP, MI or MC.** Each node represents one ramanome. The same ramanomes found in **MP**, **MI** or **MC** are linked by lines. The nodes and lines are colored based on strain.
